# The Degree Distribution of Human Brain Functional Connectivity is Generalized Pareto: A Multi-Scale Analysis

**DOI:** 10.1101/840066

**Authors:** Riccardo Zucca, Xerxes D. Arsiwalla, Hoang Le, Mikail Rubinov, Antoni Gurguí, Paul Verschure

**Affiliations:** Institute for BioEngineering of Catalonia (IBEC), Barcelona, Spain; California Institute of Technology, Pasadena, CA, USA; Department of Biomedical Engineering, Vanderbilt University, Nashville, TN, USA; Janelia Research Campus, Howard Hughes Medical Institute, Ashburn, VA, USA; Catalan Institute of Advanced Studies (ICREA), Barcelona, Spain; Barcelona Institute of Science and Technology (BIST), Barcelona, Spain

**Author notes:** Equal Contribution (Riccardo Zucca, Xerxes D. Arsiwalla).

## Abstract

Are degree distributions of human brain functional connectivity networks heavy-tailed? Initial claims based on least-square fitting suggested that brain functional connectivity networks obey power law scaling in their degree distributions. This interpretation has been challenged on methodological grounds. Subsequently, estimators based on maximum-likelihood and non-parametric tests involving surrogate data have been proposed. No clear consensus has emerged as results especially depended on data resolution. To identify the underlying topological distribution of brain functional connectivity calls for a closer examination of the relationship between resolution and statistics of model fitting. In this study, we analyze high-resolution functional magnetic resonance imaging (fMRI) data from the Human Connectome Project to assess its degree distribution across resolutions. We consider resolutions from one thousand to eighty thousand regions of interest (ROIs) and test whether they follow a heavy or short-tailed distribution. We analyze power law, exponential, truncated power law, log-normal, Weibull and generalized Pareto probability distributions. Notably, the Generalized Pareto distribution is of particular interest since it interpolates between heavy-tailed and short-tailed distributions, and it provides a handle on estimating the tail’s heaviness or shortness directly from the data. Our results show that the statistics support the short-tailed limit of the generalized Pareto distribution, rather than a power law or any other heavy-tailed distribution. Working across resolutions of the data and performing cross-model comparisons, we further establish the overall robustness of the generalized Pareto model in explaining the data. Moreover, we account for earlier ambiguities by showing that down-sampling the data systematically affects statistical results. At lower resolutions models cannot easily be differentiated on statistical grounds while their plausibility consistently increases up to an upper bound. Indeed, more power law distributions are reported at low resolutions (5K) than at higher ones (50K or 80K). However, we show that these positive identifications at low resolutions fail cross-model comparisons and that down-sampling data introduces the risk of detecting spurious heavy-tailed distributions. This dependence of the statistics of degree distributions on sampling resolution has broader implications for neuroinformatic methodology, especially, when several analyses rely on down-sampled data, for instance, due to a choice of anatomical parcellations or measurement technique. Our findings that node degrees of human brain functional networks follow a short-tailed distribution have important implications for claims of brain organization and function. Our findings do not support common simplistic representations of the brain as a generic complex system with optimally efficient architecture and function, modeled with simple growth mechanisms. Instead these findings reflect a more nuanced picture of a biological system that has been shaped by longstanding and pervasive developmental and architectural constraints, including wiring-cost constraints on the centrality architecture of individual nodes.

## Introduction

The idea that the topology of brain networks may follow power law or heavy-tailed characteristics has received a lot of attention starting with the initial discovery that some real-world networks, including social, genetic and technological networks such as the internet show power law degree distribution in scale-free networks as opposed to Poisson degree distributions in Erdos-Renyi networks^1^. Since then, power law distributions have been reported in many more instances of social, cellular and technological networks (see^2–5^ for an overview). These observations have led to the idea that most complex real-world networks may be structured, and that this structure may have arisen from simple growth mechanisms, such as preferential attachment^1^. It has also been suggested that the so-called *scale-free* property facilitates efficient communication via a small number of designated central nodes acting as hubs of information flow, as in the case of airline or transportation networks^2^. However, this interpretation has not been without controversy, and recently, it has been claimed that these cases of power law scaling might not be as prevalent as initially thought^6^ (see also^7^ for a commentary and^8^ for a counter-claim arguing that real-world scale-free networks are highly prevalent). In earlier work, challenges to the omnipresence of power laws and heavy tails have been made, but only within specific domains^9, 10^. In contrast,^6^ have analyzed power law distributions across domains taking a data-centric approach. Considering over a thousand networks from various disciplines, they conclude that scale-free networks (typically those following a power law with scaling factor close to 2) are rare in real-world data. The reason why this has only recently been realized is that confirming the existence of a statistically significant power law is a lot more demanding than previously employed heuristics of fitting data with linear least-squares on a log-log scale. As an alternative, the statistical bootstrapping method, initially developed for power law testing, was first introduced by Clauset et al. in^11^. This has since then been extended to test for other distributions and has subsequently been implemented in several studies^12–14^. However, challenges remain, both, for statistical analyses involving large data-sets (incurring a high computational cost for very large networks) and for issues concerning robustness and interpretability of results (i.e., different parcellation schemes or network representations lead to different interpretations of plausible distributions^12, 15, 16^). In this study, we address these open issues systematically, investigating the reproducibility of power laws and other heavy-tailed distributions within the specific domain of human brain functional connectivity networks constructed from resting-state functional Magnetic Resonance Imaging (rs-fMRI).

Alongside the advancement of computational and machine learning tools in data science, the neuroscience community itself has greatly benefited from adopting a network science approach. Some of the big questions in network neuroscience involve mechanisms and scaling properties of large-scale structural and functional brain networks. As in other domains of network science, there have been suggestions that the degree distribution of voxels in brain functional networks may also be scale-free or at least heavy-tailed^17–21^. These studies point to the presence of a small number of hub nodes that connect widely across the network. Other studies have suggested that functional brain networks are not scale-free, but instead are characterized by an exponentially truncated distribution^10, 22, 23^. The scaling characteristics of brain networks reflect the organization of the brain’s architecture and are therefore essential for understanding how the brain operates^24–31^ and how it responds to injury^10, 32, 33^.

The initial excitement in looking for scale-free and other heavy-tailed distributions in brain networks was due to the proposal that the dynamics of the brain might be operating at criticality^17, 18^. In this critical regime the network dynamics are scale invariant and that has been touted as a plausible mechanism for near-optimal information processing^17^. On the other hand, a definitive absence of heavy-tailed distributions in brain network topology would put this evidence for criticality at odds (see also^34^ for a critical discussion on criticality). Hence, given the current debate on power law scaling in real-world networks, a rigorous statistical analysis is required to estimate the underlying degree distribution of brain networks. Indeed, these may well not be scale-free, power law or even heavy-tailed. For these reasons, in this work, we revisit the issue of the scaling properties of human brain functional connectivity with a more detailed and specific analysis. We focus on the specific domain of brain functional networks and perform detailed checks over plausible model distributions.

The earliest studies testing for power law distributions used least-squares fitting method on log-log plots of frequency distributions of node degrees^19, 20^. This methodology, although seemingly straightforward, is statistically flawed^11^. Least-squares fitting on log-log plots gives systematically biased estimates of scaling parameters as regression lines are not valid probability distributions and do not respect normalization constraints of the associated cumulative distribution. Moreover, in addition to model testing, one also requires a statistical measure to estimate the *goodness-of-fit* of prospective degree distributions. Approaches using least-squares fitting do not consider this aspect. An alternative framework^11^ has advocated for a Maximum Likelihood Estimation (MLE) of scaling parameters and subsequent model comparisons to other distributions using samples of both, real and synthetic data. These authors have derived an analytic measure for power law models that has subsequently been extended to other distributions as well, albeit, for most of these, the derivation has to be carried through numerically.

In spite of these developments, it has been observed that distribution estimates are still dependent on a number of factors such as the way data is pre-processed (e.g. confound regression or Independent Component Analysis based de-noising procedures), how the network is constructed (e.g. using either correlations for edge weights or alternative measures of causality), how network thresholds have been set, the spatial resolution or scale of the data, and also on whether one uses region-based or voxel-based node specifications^12, 16, 23^. For instance, Hayasaka et al.^23^ have found that although degree distributions of many analyzed functional networks followed an exponentially truncated model, the higher the resolution (of the order of 15 thousand voxels), the more the distribution trends towards a power law. Hence, in response, in our data we will reanalyze degree distributions near this voxel resolution as well as those at much higher ones.

In this work, we advance an analysis which addresses the aforementioned issues. We analyze resting-state fMRI data of 10 subjects obtained from the Human Connectome Project^35^. Using maximum likelihood methods for parameter extraction, we first estimate scaling parameters for the best possible fit among all possible model distributions. Subsequently, we check the goodness-of-fit for each distribution by comparing to synthetically generated data (based on the same MLE parameters). We consider six different model distributions to cover a wide range of heavy and short-tailed distributions: power law, exponential, power law with exponential cut-off, log-normal, Weibull and generalized Pareto. In particular, the generalized Pareto distribution will be important for this study. This distribution was first introduced in^36^. Its applications include use in the analysis of extreme events, as a failure-time distribution in reliability studies^37^. In particular, it has often been employed in meteorological and geophysical studies^38–41^. What is interesting, is that this distribution interpolates between heavy-tailed and short-tailed distributions, with the power law and exponential distributions being special cases of it. This interpolation depends on a tail-parameter, which will give us a handle on estimating the tail’s heaviness or shortness directly from the data. We will consider weighted functional networks and will analyze statistics for 18 different thresholds (separately for positive as well as negative correlations) for each data-set. Furthermore, instead of choosing a fixed resolution of the data, we analyze the same data at six different voxel resolutions: 1,000, 5,000, 10,000, 20,000, 50,000 and 80,000 voxels. These resolutions are obtained by down-sampling the original data at *∼*80K voxels to lower resolutions.

Importantly, our results show that the topology of human brain functional connectivity networks follow a short-tailed distribution. Additionally, we demonstrate that down-sampling data introduces the risk of detecting spurious heavy-tailed distributions that fail cross-model comparisons. This dependence of statistics on data resolution has broader implications for neuroinformatic methodologies and analyses, especially, when these analyses rely on down-sampled data (for instance, into anatomical parcellations). Our findings that node degrees of brain functional networks follow a short-tailed distribution have important implications for prospective brain architectures, including realistic biological constraints such as wiring cost and aging of nodes.

## Methods

### Participants, Imaging Data and Network Extraction

We analyzed high-quality, high-resolution resting-state fMRI scans of 10 subjects (age range: 26 to 35, 16.7% male) obtained from the Human Connectome Project (HCP, Q1 data-set, released by the WU-Minn HCP consortium in March 2013^35^). Individual rs-fMRI data were acquired for *∼*15 minutes providing a total of *∼*80,000 gray-ordinates time-series of 1,200 time points each (min-max range 67,709 - 86,332). Before analyzing it, each data-set has been transformed into a series of graphs. Data preprocessing involved ICA de-noising of the time-series in order to remove artefacts. A schematic illustration of the overall procedure used to build each of the networks is provided in Fig. 1A. Building and visualizing functional networks was performed using the BrainX^3^ platform^33, 42–44^.

**Figure 1.**
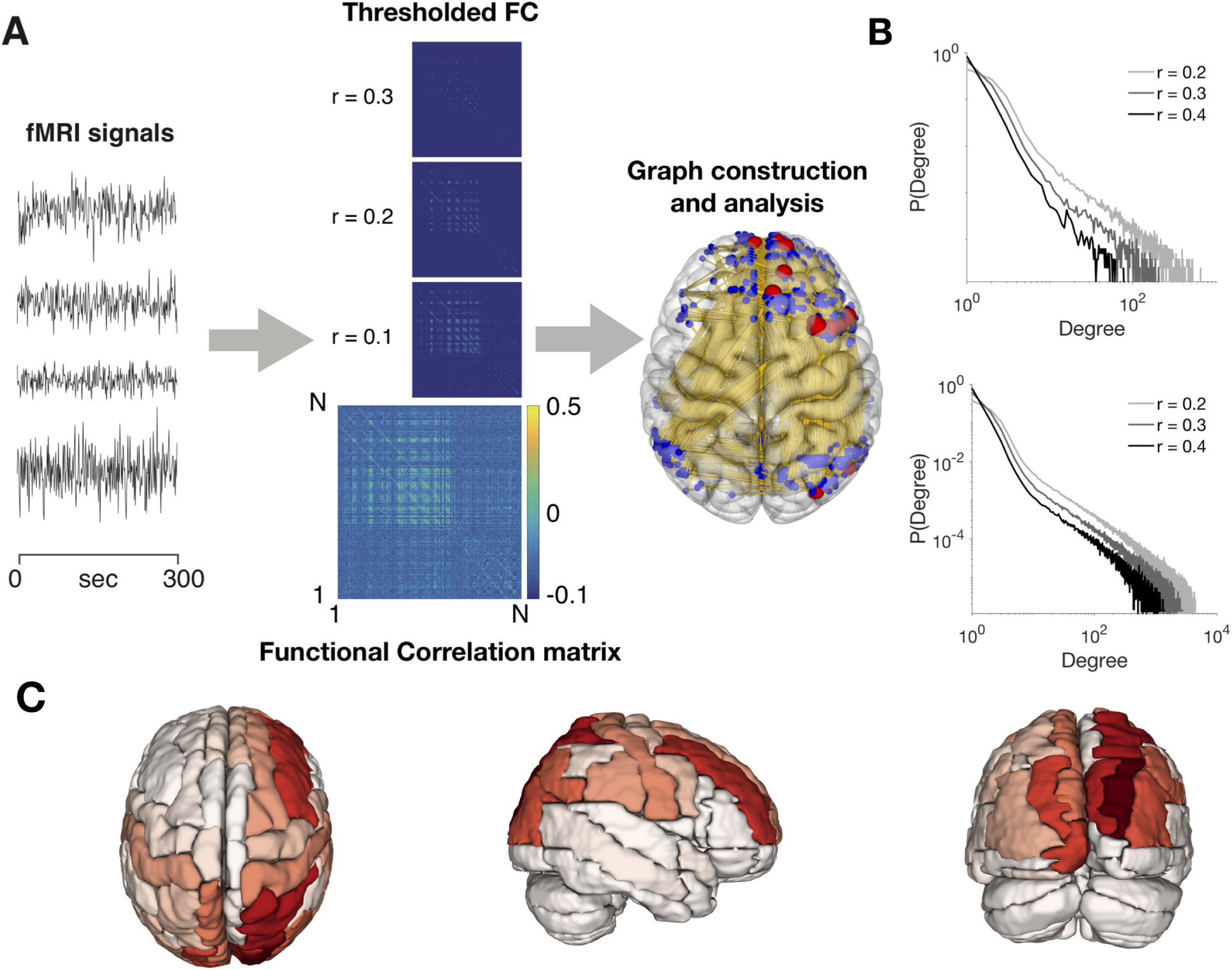
**A.** Overview of the processing steps used to generate graph-based brain connectivity functional networks. Five different parcellation schemes were generated which divided the original *∼* 80K brain data into 1K, 5K, 10K, 20K, 50K regions-of-interest (ROIs). For each node pair, temporal correlation was calculated from the fMRI signals to generate a Functional Connectivity (FC) matrix for each subject. The edges’ distribution of the resulting individual weighted functional networks is then examined for a range of different thresholds (examples are given for thresholds equal to 0.1, 0.2 and 0.3) from which distinct graph structures can be defined. **B.** An example of degree distributions for three different values of the FC threshold for a representative data-set (top) and the average over the 10 data-sets included in the study (bottom). **C.** Region-wise group average for the 80K resolution of the top twenty hubs mapped on the Automated Anatomical Labeling (AAL) volume atlas^46^. Across subjects the highest degree connectivity is observed across the fronto-parietal-occipital areas. Darker colors denote regions belonging to a larger number of subject data-sets.

For all the subjects, we consider six different network resolutions where the original data-set of *∼*80,000 regions of interest (ROIs) is further down-sampled at five different resolutions *∼*1,000, *∼*5,000, *∼*10,000, *∼*20,000 and *∼*50,000 regions of interest by averaging the time-series of neighbouring gray-ordinates within a cube of 13, 7, 5, 4, 2.5 mm^3^, respectively (see supplementary table 4 for the exact number of ROIs for each data-set).

We build a *N × N* functional connectivity matrix for each data-set by calculating the Pearson’s correlation coefficient between each possible pair of ROIs^45^, where *N* corresponds to the number of nodes in the network, which is symmetric by construction and with self-connections set to zero. Our analysis took into account the weighted degree distributions of the data. We examined the full range of positive as well as negative correlation thresholds. To obtain weighted un-directed adjacency matrices, we threshold each functional matrix at 18 different levels (*R*) in the range *−*0.7 to 0.8, at intervals of 0.1. Outside this range, the functional matrices become too sparse for meaningful statistics. For positive thresholds (*r >* 0), the weighted adjacency matrix is obtained by keeping all the values above threshold while all entries below threshold are set to zero. For negative correlations, a threshold sets an upper bound. All correlation strengths more negative than the threshold are maintained in the adjacency matrix, whereas correlations above the threshold are set to zero. In a weighted network, the weighted degree of a node corresponds to the sum of all weighted edges connected to that node. Figure 1B depicts the degree distributions of extracted networks across three different thresholds (r *>* 0.2, 0.3 and 0.4) for a representative subject and the averaged degree distributions over all ten data-sets. The region-wise group average for the original dataset of the top twenty leading degree nodes mapped on the Automated Anatomical Labeling (AAL) volume atlas^46^ is illustrated in Figure 1C.

### Fitting Parametric Models to Weighted Degree Networks

For every generated network, the vector of degrees **x** = [*x*_1_*, x*_2_*, …, x_n_*] is sorted in ascending order for each correlation threshold. Fitting parametric models to these degree distributions follows the statistical bootstrapping approach outlined in^11^. This method uses *Maximum Likelihood Estimation* (MLE) to determine model parameters, followed by the Kolmogorov-Smirnov (KS) statistic to estimate the tail of the distribution corresponding to that model. For example, in the case of power law models, MLE is used to estimate the scaling parameter *α* providing the best possible fit for a hypothetical power law distribution *P*(*x*) *∼ Cx^−α^* for the tail of the observed data in the range *x_i_* to *x_n_*. Next, we determine the KS statistic for this power law distribution with respect to *x_i_*. Out of all possible *x_i_* from the data, the one with the smallest KS statistic corresponds to the lower bound *x_min_* for power law behavior in the data. For other models (listed in table 1), the same procedure is repeated to estimate their respective parameters.

**Table 1.** The six models used for the analysis of the degree distribution.

| Distribution model | Probability density function (PDF) |
| --- | --- |
| Power law | $x^{-\alpha}$ |
| Exponential | $e^{-\lambda x}$ |
| Power law with exponential cutoff | $x^\alpha e^{\beta x}$ |
| Weibull | $x^{(\beta-1)} e^{-\lambda x^\beta}$ |
| Log-Normal | $\frac{1}{x} e^{-\frac{(\log(x)-\mu)^2}{2\sigma^2}}$ |
| Generalized Pareto | $(1 + k \frac{x-x_{min}}{\sigma})^{-1-\frac{1}{k}}$ |

To verify whether the observed KS statistic indeed provides a good fit for the data, we then generate and fit 1000 synthetic data-sets from a true model distribution using the parameters determined from MLE and the bound *x_min_* as the one estimated for the best fit of the empirical data. We then fit each synthetic data-set by calculating the KS statistic relative to its original model. From that, we calculate an empirical *p*-value as the fraction of the times the empirical distribution shows a smaller value of the KS statistic as compared to the synthetically generated ones. If the obtained *p*-value is below a significant threshold, *p* ⩽ 0.1^11^, the model hypothesis can be ruled out as a non-plausible explanation of the data. Furthermore, we impose an additional constraint that the tail size of a plausible distribution contains at least fifty nodes, to avoid those cases where the *p*-value may be high, but the tail is extremely sparse. Note, however, that large *p*-values by themselves do not guarantee that the given model is the best. One still has to perform cross-model comparisons with other plausible distributions (listed in table 1).

All the analyses were performed in Matlab (Mathworks Inc., USA) using the methods from^11^. Further, for testing the alternative models, we adapted the framework provided in^11^ to include the competing hypothesis. For each subject, we analyzed thresholds in the range *−*0.7 to 0.8, with 0.1 increments. The parametric *goodness-of-fit* test was conducted over 1,000 repetitions, ensuring precision of *p*-value up to two decimal digits. Fittings to power law distribution for the 10K resolution were also computed using the *Powerlaw* Python package from^14^ to verify the consistency of the procedure used here.

## Results

In order to estimate degree distributions of human brain functional networks, we analyzed high-resolution fMRI data from the Human Connectome Project at varying resolutions from one thousand to 80 thousand regions of interest (ROIs) and tested whether they follow heavy or short-tailed distributions considering the power law, exponential, power law with an exponential cutoff, log-normal, Weibull and generalized Pareto distributions. We tested each of the above-mentioned statistical models for 18 different functional connectivity thresholds in each of the ten subjects and across all resolutions of the data (1K, 5K, 10K, 20K, 50K and 80K).

Our analysis revealed that across all resolutions of the data the statistical plausibility of the power law model is consistently weaker than other models (with the exception of the Weibull distribution; Figure 2 left panel and Table 2). Indeed, it is the generalized Pareto model that consistently dominates the statistics at every resolution (mean proportion of statistically plausible fits across resolutions: 49.98 *±*4.81). Another trend we observe is the increase in the proportion of plausible model fits as the data is down-sampled (Figure 2B). In particular, this increase is strictly monotonous for all models from a resolution of 50K to 10K. As we shall see, this is because at lower resolutions multiple models become simultaneously statistically significant (Figure 2B).

**Figure 2.**
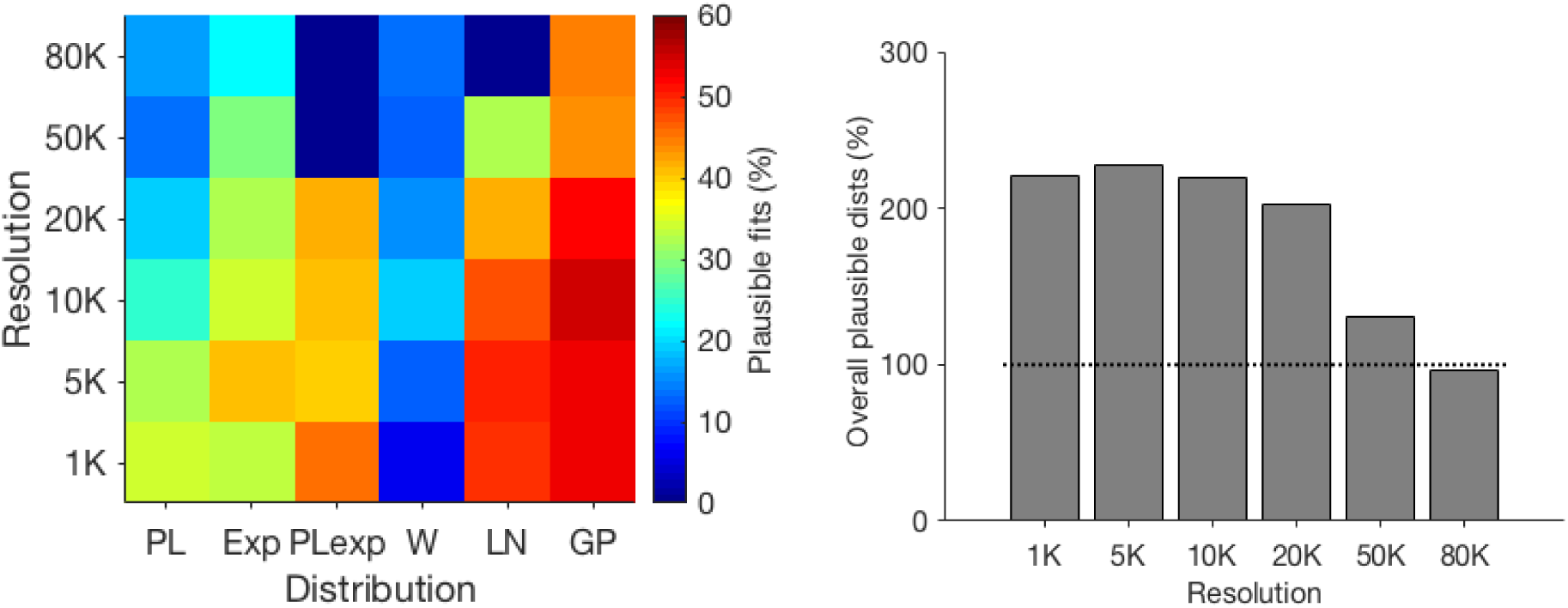
Overall results of model fitting. Left: Proportion of fitted distributions that are statistically significant. Across resolutions, the generalized Pareto model is consistently outperforming the other candidate models. Right. Proportion of models that are a plausible explanation of the data across resolutions. For lower resolutions, multiple models become simultaneously plausible. A fit is considered plausible if its p-value is equal or larger than 0.1 and the tail of the distribution contains more than 50 nodes.

**Table 2.** Proportion of fitted distributions that are a statistically plausible explanation of the data with the goodness-of-fit test larger or equal to 0.1. Legend: PL: power law, Exp: exponential, LNOrm: log-normal, Wei: Weibull, GP: Generalized Pareto, PLexp: Power law with exponential cutoff, na: numerical analysis did not converge to a stable solution.

|  | PL (%) | Exp (%) | PLexp (%) | LNorm (%) | Wei (%) | GP (%) |
| --- | --- | --- | --- | --- | --- | --- |
| 80K | 16.7 | 21.7 | na | na | 13.3 | 44.4 |
| 50K | 13.3 | 29.4 | na | 32.2 | 12.2 | 43.3 |
| 20K | 19.4 | 32.2 | 41.7 | 41.7 | 15.0 | 52.2 |
| 10K | 24.4 | 34.4 | 40.6 | 47.2 | 18.9 | 54.4 |
| 5K | 32.2 | 40.6 | 39.4 | 50.0 | 12.2 | 52.8 |
| 1K | 34.4 | 33.3 | 45.0 | 48.9 | 6.1 | 52.8 |

After model fitting, when multiple models became simultaneously plausible, we performed log-likelihood ratio (LLR) tests between pairs of models to determine the most plausible one. The log-likelihood tests have been done at each FC threshold, for each subject, and each resolution.

Once again, percentages of the number of times a given model outperforms a competing model in pair-wise comparisons indicates a clear dominance of the generalized Pareto model compared to all other distributions and this superiority is consistent across all resolutions (Figure 3, Table 3 and supplementary Tables 38 to 43). In contrast, the power law distribution turns out to be the weakest (statistically) model in log-likelihood ratio tests, at every resolution. The percentages of log-likelihood ratio outcomes are fairly robust across resolutions with the exception of the highest resolution, where, as noted earlier, there is a lower number of multiple comparisons. What is also noteworthy is the number of ‘inconclusive’ cases (Table 3 and Figure 3). These indicate instances when it was statistically impossible to discern between multiple models. These occurrences are the lowest at the highest resolution and rise systematically with every decreasing data resolution. What we find is that coarse-graining the original data by half already leads to a threefold increase in the number of inconclusive statistical tests as compared to the highest resolution. power law model, by definition, is heavy-tailed, whereas the exponential is short-tailed. The generalized Pareto, log-normal and Weibull all have parameters that explicitly determine the shape of the tail (*k*, *σ* and *β* respectively). The generalized Pareto model is of particular interest here as it interpolates between heavy-tailed and short-tailed distributions. A large positive *k* value indicates the presence of a fat-tail, whereas a small or negative value points to the opposite. Given that the generalized Pareto model statistically outperforms all other models in our analysis, across thresholds and resolutions, we examined the values of its *k* parameter for those instances where the p-value of the model is greater than 0.1 and tail size is higher than 50 (Figure 4A). Overall, including both positive and negative thresholds, we find that *k* is close to zero, approaching an exponential distribution. When considered separately, for positive thresholds *k* assumes negative values, implying a short tail (Figure 5B), whereas it becomes positive for negative thresholds. Moreover, these observations hold across all resolutions (Figure 5C).

**Figure 3.**
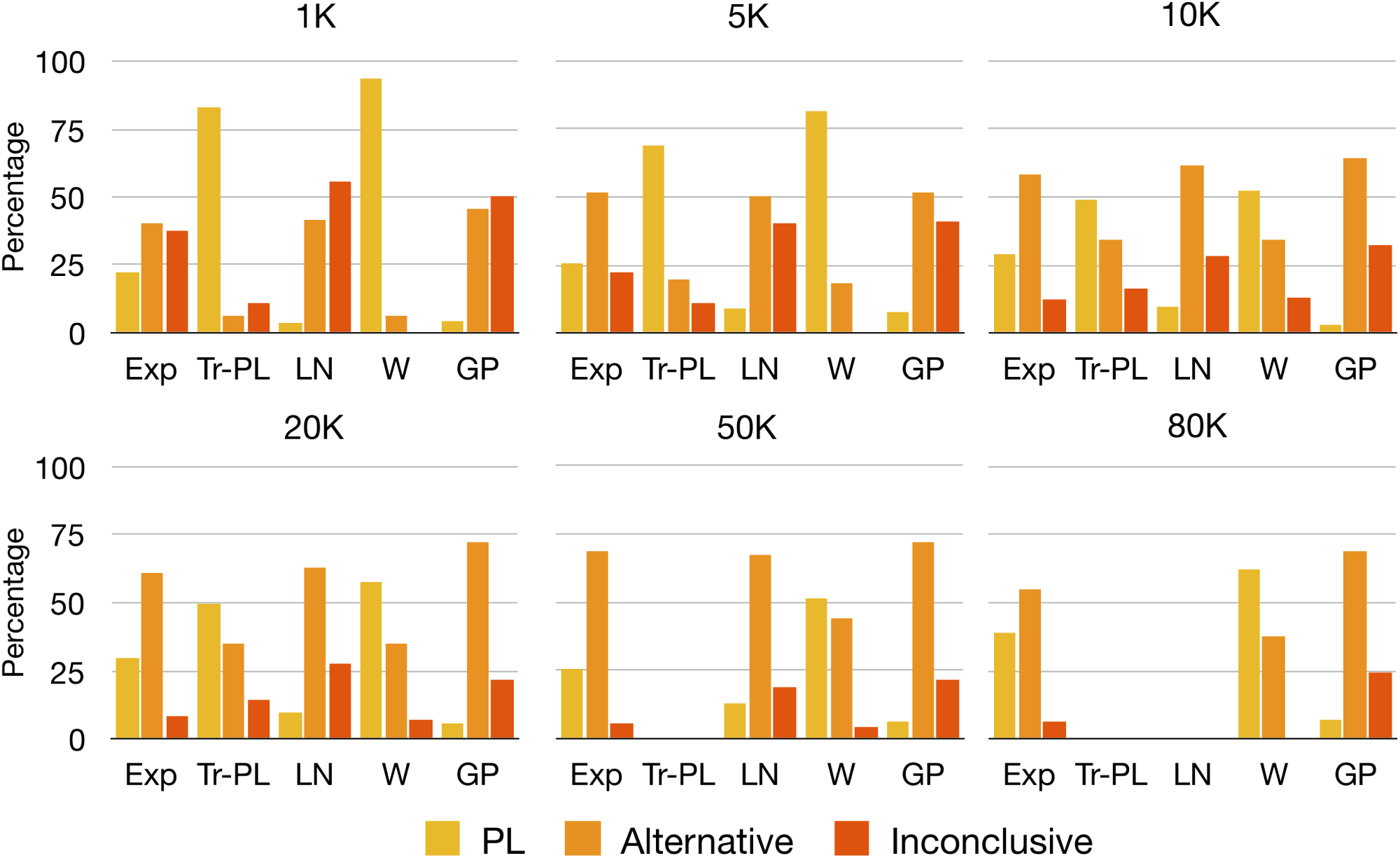
Log-likelihood ratio test results from comparing the best fit for alternative distributions with the best fit power law distribution. We show the percentage of times a power law model (PL), the alternative model (Alternative) or neither (Inconclusive) was favored.

**Figure 4.**
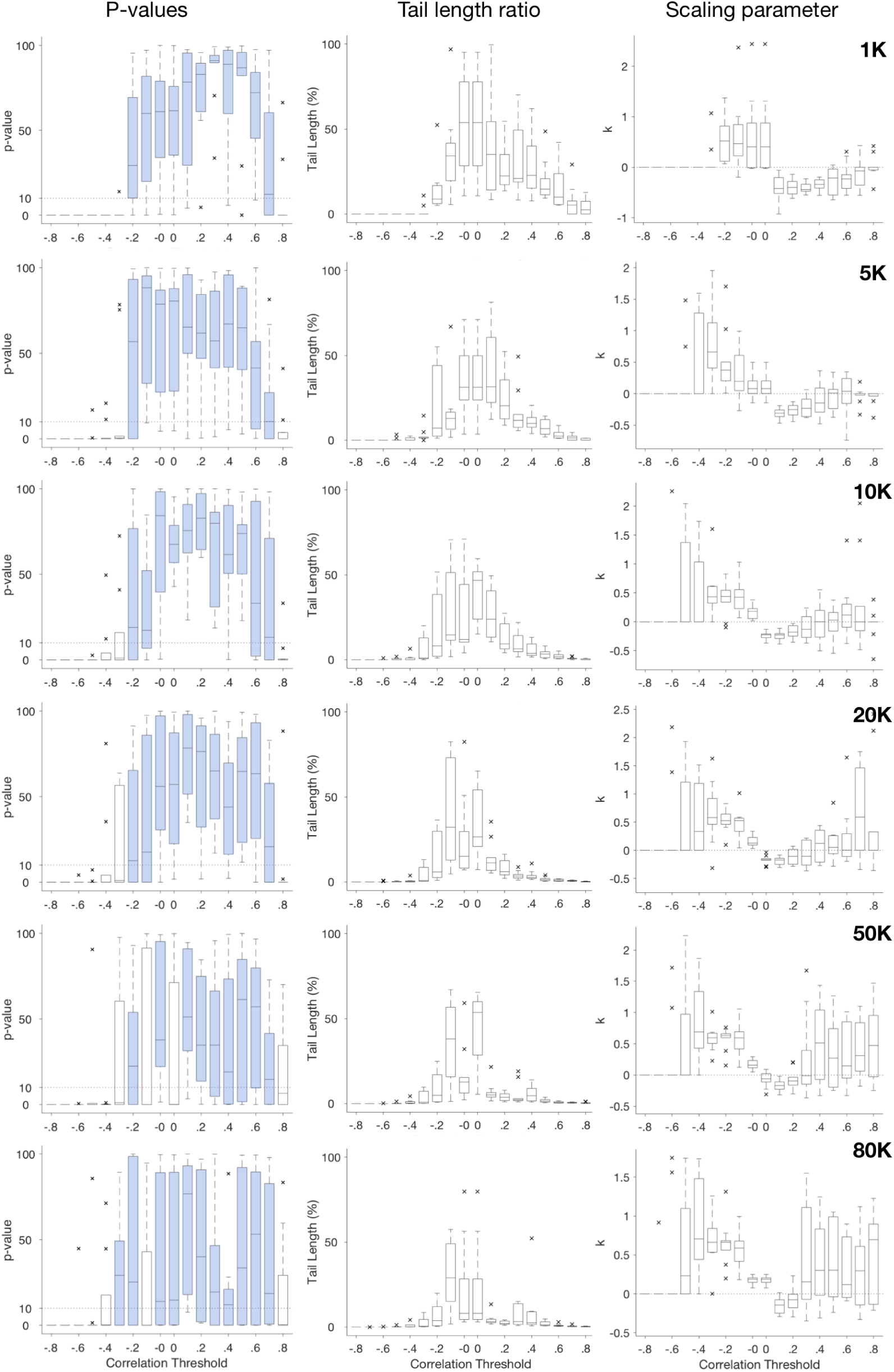
Statistics of the generalized Pareto model across resolutions and thresholds. Left: population averaged goodness-of-fit tests. Center: percentage of the tail of the distribution explained by the model across different thresholds. Horizontal dashed lines in the box-plots indicate the acceptance criteria for a model to be considered plausible (p-value*>* 10). The central mark is the median, the edges of the boxes are the 25*^th^* and 75*^th^* percentiles. Right: Estimated *k* parameter as a function of threshold.

**Figure 5.**
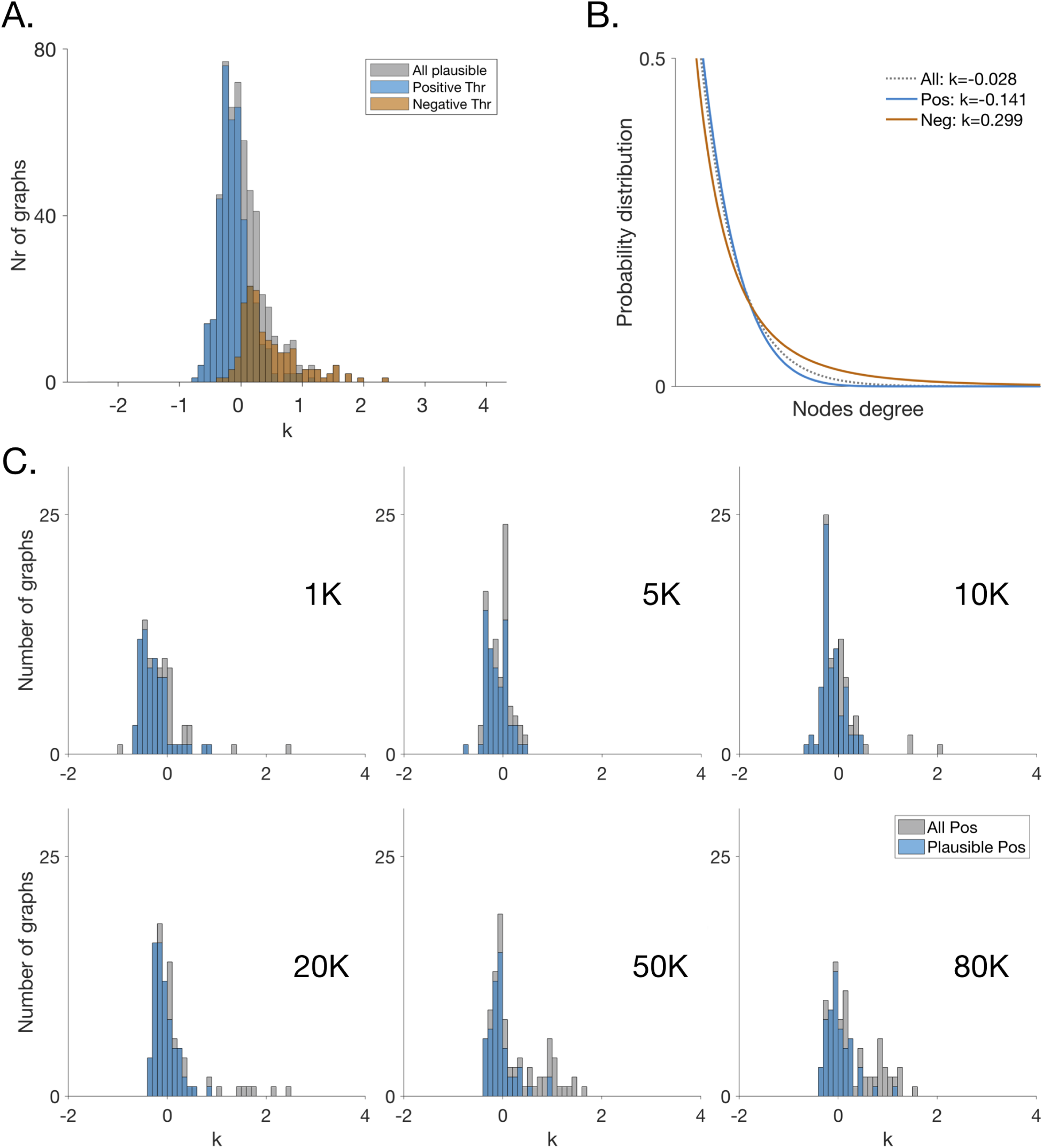
The degree distributions of functional connectivity networks tend towards the shorter limit of the generalized Pareto distribution. A: Overall distribution of the generalized Pareto estimated *k* parameter values for those distributions that are a statistically plausible fit of the data (p-value>0.1 with a minimum tail length of 50 nodes). Large positive *k* values indicate the presence of a heavy-tail, whereas small or negative values point to a suppressed tail. B: Example of short- and heavy-tailed generalized Pareto distributions as a function of the scaling parameter *k*. C. Distributions of the generalized Pareto estimated *k* parameter values for the positive thresholds across resolutions. Gray bars corresponds to all possible fits whereas blue bars corresponds to statistically plausible fits that (p-value>0.1 with a minimum tail length of 50 nodes)

**Table 3.** Overall Likelihood ratio test results of comparing the alternative distributions at the different resolutions. The data express the number of times a model is a plausible explanation of the data. In parenthesis the proportion with respect to the total number of comparisons made.

| Distribution | 1K | 5K | 10K | 20K | 50K | 80K |
| --- | --- | --- | --- | --- | --- | --- |
| Power law | 141 (11.6) | 148 (11.3) | 114 (8.9) | 100 (8.6) | 56 (7.4) | 61 (13.9) |
| Exponential | 162 (13.3) | 189 (14.4) | 167 (13.1) | 164 (14.2) | 140 (18.6) | 80 (18.3) |
| Truncated PL | 85 (7.0) | 112 (6.6) | 136 (10.6) | 115 (9.9) | - (-) | - (-) |
| Log normal | 230 (18.9) | 229 (17.5) | 219 (17.1) | 198 (17.1) | 141 (18.7) | - (-) |
| Weibull | 20 (1.6) | 54 (4.1) | 99 (7.7) | 89 (7.7) | 62 (8.2) | 53 (12.1) |
| Gen Pareto | 265 (21.8) | 260 (19.9) | 276 (21.6) | 282 (24.4) | 231 (30.7) | 191 (43.6) |
| Inconclusive | 312 (25.7) | 316 (24.1) | 267 (20.9) | 207 (17.9) | 122 (16.2) | 53 (12.1) |
| Comparisons | 1215 | 1308 | 1278 | 1155 | 752 | 438 |

Our analysis also reveals the shape of the tail of degree distributions of functional networks allowing to answer the question whether they are heavy-tailed or short-tailed. This question can be addressed by looking at the most plausible statistical model at each threshold and examining the values of the tail parameters of the model – when such parameters explicitly exist. The Taken together, the main conclusions of our analysis is that:

i. Degree distributions of brain functional connectivity networks obtained from fMRI recordings rarely follow a power law scaling. Instead, the generalized Pareto distribution provides the best statistical explanation of the graph of the functional connectivity network of the human brain.
ii. The results pertaining to model comparisons remain robust at all resolutions of the data from the very high resolution of 80K voxels up to the low resolution of 1K.
iii. The degree distributions of these networks are rarely heavy-tailed. Instead, the trend is towards short-tailed distributions. We see this from values of the shape parameter *k* of the generalized Pareto distribution as well as *σ* of the log-normal distribution.
iv. Data down-sampling systematically affects all statistical tests. Namely, the statistical plausibility of every model increases upon down-sampling. Lowering the resolution of the data makes it harder to statistically discern between models, even though the actual tail lengths only decrease moderately as we lower resolutions as seen in figure 4). Multiple models become simultaneously plausible at low resolutions.

## Discussion

We have examined degree distributions of human brain functional networks constructed from high-resolution resting-state fMRI data to clarify contrasting claims made in the literature concerning the nature of their underlying graph. The main conclusion of our analysis is that these networks are short-tailed, following the generalized Pareto distribution. While several alternatives to the power law and other heavy-tailed models have been extensively discussed, the generalized Pareto model has surprisingly received little attention outside of meteorology and geophysics^38–41^. What is remarkable, is that this distribution interpolates between heavy-tailed and short-tailed distributions, with the power law and exponential distributions being special cases of it. This interpolation depends on a tail-parameter, which gives one a handle on estimating the tail’s heaviness or shortness directly from the data. Here, we have found that the generalized Pareto distribution happens to outperform all other distributions, at least, within the domain of human brain functional networks.

Overall, our results indicate that the statistics do not support a heavy-tailed network topology for node degree distributions of human brain functional networks. The heavy-tail hypothesis, including the power law is firmly rejected in the majority of the thresholds we examined. Instead, it is the generalized Pareto distribution that is consistently preferable to competing models for most of the examined thresholds. We also tested for other models commonly discussed in the literature, such as the exponential, log-normal, Weibull and power law with exponential cut-off (truncated power law). Overall, we find that the generalized Pareto model outperforms all others across resolutions. These results suggest that after taking into account continuously weighted networks at each threshold (rather than binary networks), the dynamics of brain functional networks might not be governed by as many ultra-high degree hubs as a typical heavy-tailed network.

For completeness, let us also mention how our results are affected by specific parameter settings. Note that the generalized Pareto, Weibull as well as the exponentially truncated power law are all defined by three parameters (scale or normalization factor, shape factor, and tail parameter), whereas the power law, exponential and log-normal are defined using only two parameters (scale/normalization factor and tail parameter). One may ask whether the improved statistical significance is merely the result of adding an extra parameter to the model. One can see that this is not the case as the Weibull and the truncated power law do not systematically outperform any of the two-parameter distributions. It is only the generalized Pareto within its short-tail limit that best characterizes the shape of the tail in the data.

What does the above observation mean for the heavy-tailed hypothesis (sometimes also referred to as the fat-tailed hypothesis) in relation to brain functional networks? Our results on model tail parameters suggest that human brain functional networks have a short tail, rather than a heavy or fat tail as observed in the estimated *k* parameter values of the generalized Pareto model. Since this model interpolates between heavy-tailed and short-tailed distributions it includes both power law and exponential distributions as special cases. Most *k* values (considering only models passing plausibility criteria) of networks studied here point to a short tail, in many instances, even shorter than the exponential. In other words, these models have even fewer ultra high degree nodes than what would be expected for a random graph. In terms of the implications that this might have, let us make the following remarks. Firstly, this network design could be explained as an outcome of pervasive developmental and architectural constraints, including wiring-cost constraints, which prevent the emergence of long-range hubs, under the assumption that long-distance functional connectivity connections correspond to long-distance anatomical connections (a hypothesis that can be experimentally tested in future studies). Secondly, in a modeling study carried out in^47^, the authors showed how constraints to a preferential attachment growth model limit the shape of the resulting tail. More specifically, this study showed that when either the cost of adding new edges to existing vertices increases sufficiently, or that a significant number of vertices become inactive due to aging processes, then the network topology inevitably settles to a short-tailed distribution, in spite of a preferential attachment growth model. Given that such constraints on edge costs and vertex aging are reasonable for brain networks, this study lends credence to our conclusion that the analyzed functional connectivity networks may be short-tailed. And more generally, our findings do not support common simplistic representations of the brain as a generic complex system with optimally efficient architecture and function, modeled with simple growth mechanisms. Instead these findings reflect a more nuanced picture of a biological system that has been shaped by longstanding and pervasive developmental and architectural constraints, including wiring-cost constraints on the centrality architecture of individual nodes^48–50^.

An important observation emerging from our study concerns the effect that down-sampling of data has on statistical models. We found that down-sampling systematically affects all statistical tests. One might think that down-sampling smooths out variations in the data leading to more robust statistics. The opposite turns out to be the case. Intrinsic variability present in data at higher resolutions helps differentiate between competing models, enabling greater interpretability of observed results. On the other hand, we found that down-sampling the original data by half already leads to an increase in inconclusive model comparisons by more than three times the original. This is because the statistical plausibility of every model systematically increases, even though the actual tail lengths only decrease moderately as we lower resolutions. The result is that multiple models simultaneously pass plausibility criteria, making it harder to discern between them. Similar to data over-fitting, there are not sufficient features in the down-sampled data to distinguish between models. Of course, at very low resolutions, this effect breaks down as the tail of the degree distributions in the data by necessity begins to get sparse. Note that even if the data at each resolution were to be explained by a different model, the point here is that unless cross-model comparisons show statistical discernibility, those results have to be interpreted with caution. Thus, at low resolutions of the data, one does see more power laws than at higher resolutions, but those fail cross-model comparisons. This point is particularly relevant for studies where one routinely down-samples functional data into anatomical parcellations, for instance, when comparing fMRI data to various neuro-computational models^51, 52^. Even though the focus of our work here concerned the identification of the underlying topological distribution of human functional connectivity and the reproducibility of power laws in human fMRI data, the down-sampling effects we have reported bear significance for the broader discussion of reproducibility of scientific results^53, 54^. Many of the problems associated with reproducibility have been attributed to flawed methodology^54^. Within the narrow domain of the problem we have addressed here, methodological rigour turns out to be extremely important to verify robustness and interpretability of results. Statistical significance is a necessary condition, but, by itself, is not sufficient. Goodness-of-fit tests and discernibility in cross-model comparisons turn out to be methodologically crucial for reproducible science.

Finally, how does our study address the on-going debate on the abundance (or universality) of power law networks?^6, 7^ As proposed in^55^, “knowledge of whether or not a distribution is heavy-tailed is far more important than whether it can be fit using a power law”. Extending this philosophy, an empirical detection of a statistical distribution can be insightful either when it brings us closer to understanding underlying organizational principles or results from one. In the current study, evidence favoring a short-tailed, rather than a heavy-tailed degree distribution suggests constraints on the topological organization of brain networks. There have also been criticisms against the conclusions of statistical tests applied to real-world data, claiming that such tests will always discriminate against power laws because strictly speaking, power laws are only to be found in the infinite size limit of growth models as preferential attachment. To counter this claim, we point to studies where the same statistical methods used here have also been able to rigorously establish power law behaviors in temporal dynamics of localized fMRI signals as well as brain electric field potentials, without having to resort to asymptotic limits^56, 57^. In those studies, power law scaling underlies heavy-range temporal correlations. Besides that, as pointed out in^55^, under certain conditions, power laws also arise from mixing multiple heavy-tailed distributions (as a special case of the central limit theorem). Hence, the infinite limit argument cannot be used every time a statistical test fails to show a power law in the data. In systems where such a limit is physically meaningful and can be justified using a growth model, this argument would have been plausible. However, brain functional networks are finite-sized and follow more complicated growth patterns. Nonetheless, the network sizes we have examined here are much larger than those considered in previous studies of functional networks. At the highest resolution, there is no trend towards a power law, quite the opposite. At higher resolutions of the data, the trend moves away from a power law, and the data shows evidence for short tails. In this case, we would conclude that observed finite size effects provide useful indicators to probe underlying informational and organizational principles of brain function. Another suggestion, made recently in^8^, is to test for “noisy power laws”, that are modulated by slowly varying functions which approach a constant in the large size limit. This point is well taken. However, in our analysis, we have considered distributions that interpolate between heavy-tailed and short-tailed distributions, including possible modulations of power laws. Once again, we find that the data points in the direction of short-tails. Even more recently,^58^ have suggested that power law tests can be affected by correlations present in the data, which may lead to their false rejections. To resolve this, they proposed a method based on shuffling and under-sampling the data to account for correlations. We have addressed this issue in our analysis by way of down-sampling across resolutions. It has been noted in^12^ that down-sampling is yet another way to control for the effects of local correlations. Indeed, at lower resolutions, more power laws pass compared to higher ones. Nevertheless, in our data, we found that the generalized Pareto model consistently outperforms the power law model at all resolutions. In summary, for many real-world problems, including brain dynamics, finite size effects are not merely statistical fluctuations about a “true” underlying theory, but signatures of new systems-level principles. Therefore, for future work, the development of rigorous computational methods for the analysis of order parameters and non-equilibrium effects in real-world networks will prove valuable for the network science community.

## Acknowledgements

This work has been supported by the European Research Council grant agreement no. 341196 cDAC to P. Verschure. The data used in this study was made available by the Human Connectome Project, WU-Minn Consortium (Principal Investigators: David Van Essen and Kamil Ugurbil; 1U54MH091657) funded by the 16 NIH Institutes and Centers that support the NIH Blueprint for Neuroscience Research; and by the McDonnell Center for Systems Neuroscience at Washington University.

## Supplementary material

**Table 4.** List of datasets used for the analysis. Each column indicates the number of nodes at the six different resolutions. Data belong to the Q1 data released by the WU-Minn Human Connectome Project consortium in March 2013^35^

| ID | 1K | 5K | 10K | 20K | 50K | 80K |
| --- | --- | --- | --- | --- | --- | --- |
| 100307 | 1012 | 5026 | 10070 | 20151 | 50005 | 79547 |
| 103414 | 1006 | 5012 | 10072 | 20003 | 50089 | 82503 |
| 105115 | 1006 | 5009 | 10041 | 20086 | 50127 | 84857 |
| 110411 | 1006 | 5041 | 10039 | 20027 | 50376 | 67709 |
| 111312 | 1011 | 5000 | 10021 | 20115 | 50015 | 72671 |
| 113619 | 1009 | 5062 | 10017 | 20081 | 50006 | 76703 |
| 115320 | 1002 | 5004 | 10074 | 20104 | 50111 | 76802 |
| 117122 | 1097 | 5000 | 10020 | 20064 | 50157 | 76460 |
| 118730 | 1004 | 5010 | 10024 | 20079 | 50241 | 86332 |
| 118932 | 993 | 5030 | 10008 | 20040 | 50114 | 85292 |

**Figure 6.**
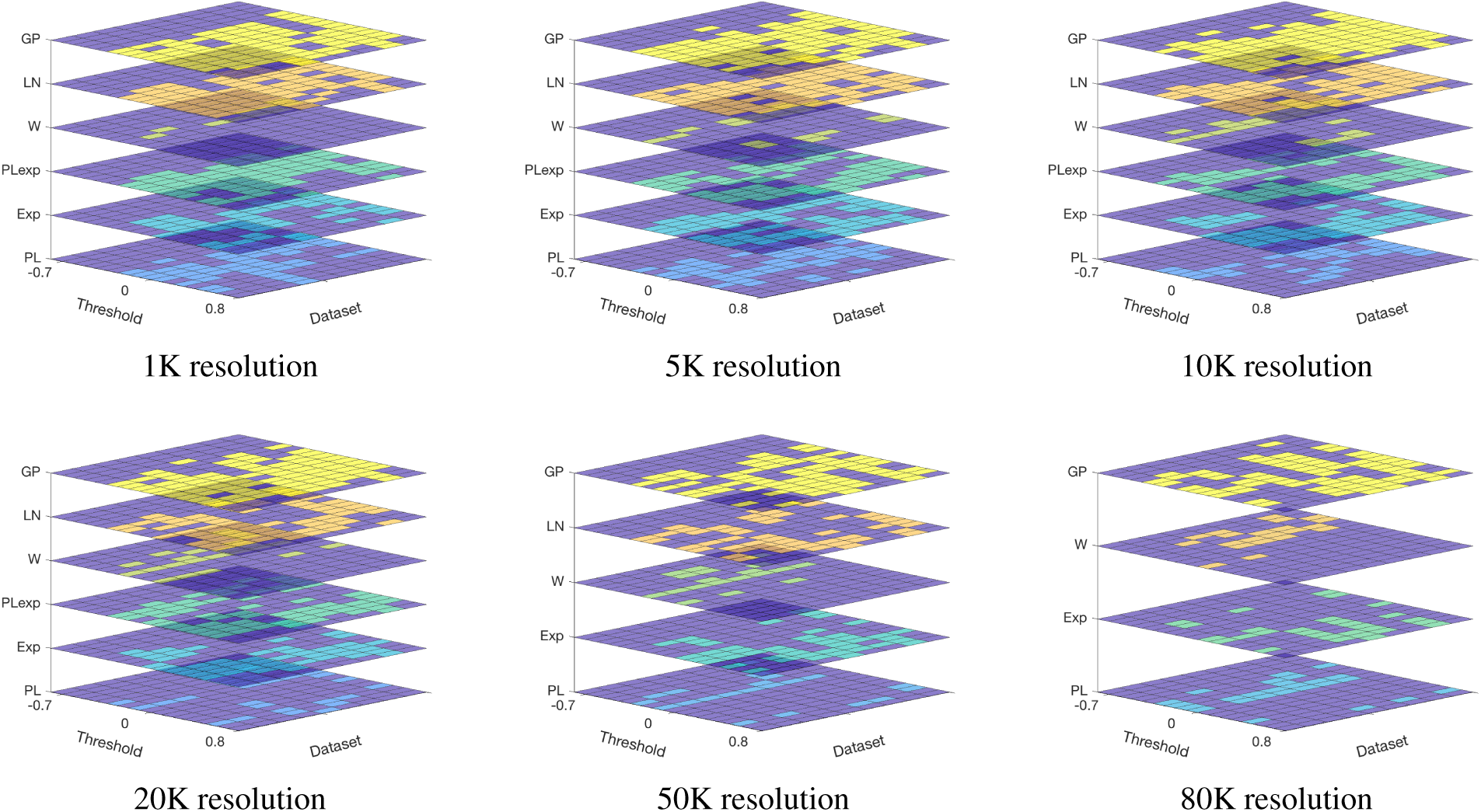
Overall distribution of the plausible fits (p*>*0.1 and tail *>*50) for the different examined thresholds and resolutions.

**Figure 7.**
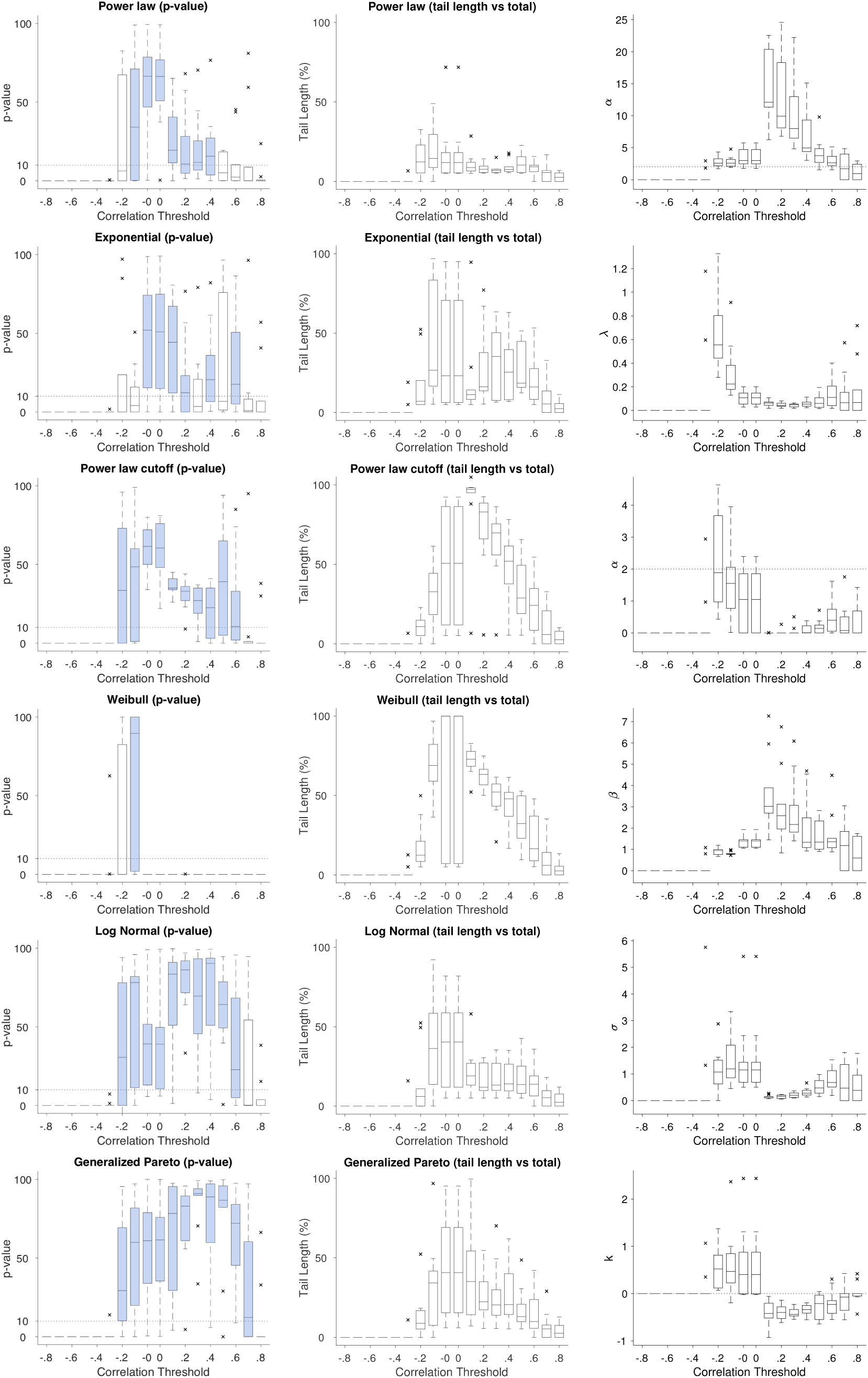
1K data set. Left: population averaged goodness-of-fit tests. Center: percentage of the tail of the distribution explained by the model (center) across different thresholds for each of the six distributions. Horizontal dashed lines in the box-plots indicate the acceptance criteria for a model to be considered plausible (p-value*>* 10).The central mark is the median, the edges of the boxes are the 25*^th^* and 75*^th^* percentiles. Cross marks correspond to outliers. Right: Estimated scaling parameters for the different candidate distributions as a function of threshold.

**Figure 8.**
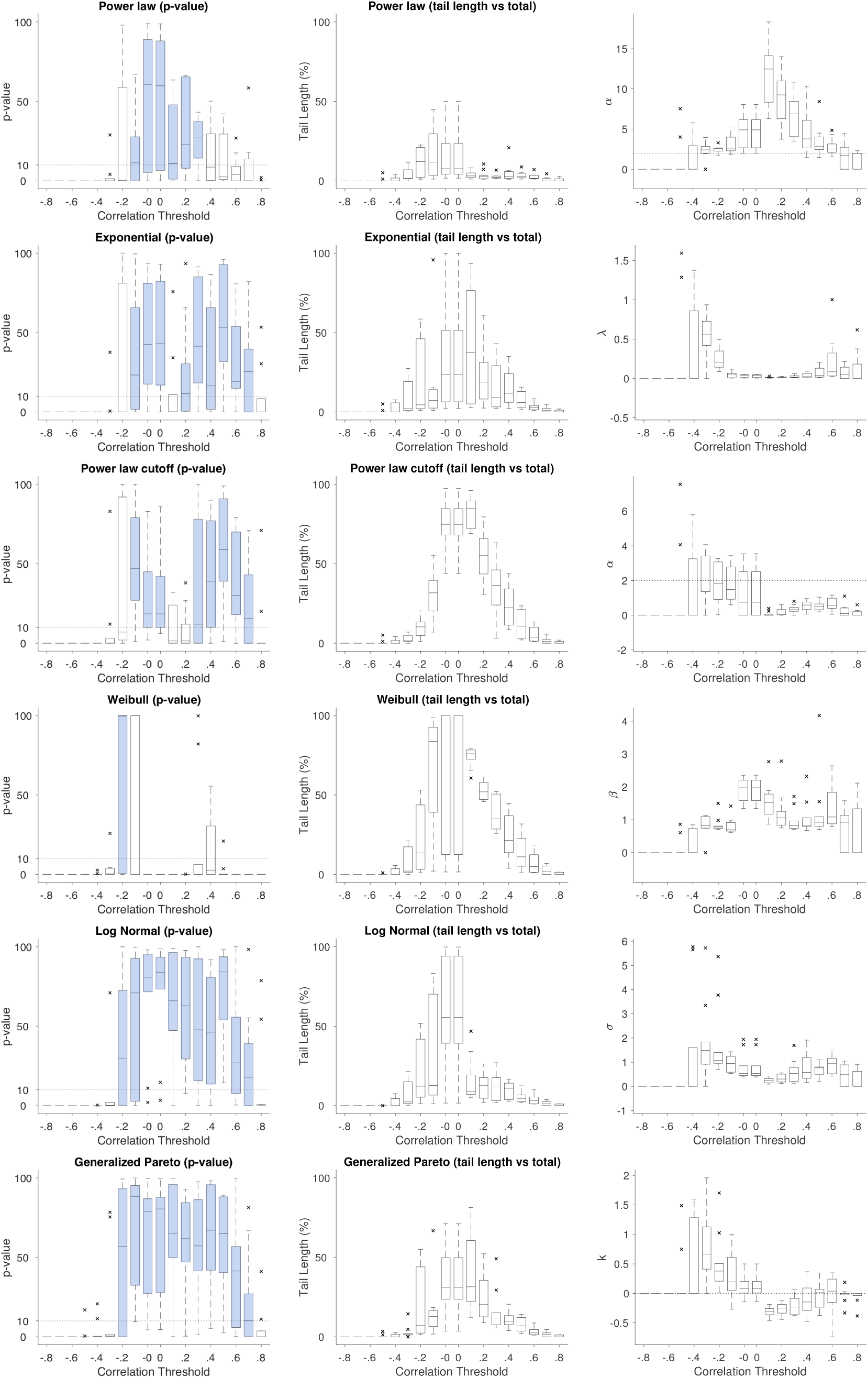
5K data set. Left: population averaged goodness-of-fit tests. Center: percentage of the tail of the distribution explained by the model (center) across different thresholds for each of the six distributions. Horizontal dashed lines in the box-plots indicate the acceptance criteria for a model to be considered plausible (p-value*>* 10).The central mark is the median, the edges of the boxes are the 25*^th^* and 75*^th^* percentiles. Cross marks correspond to outliers. Right: Estimated scaling parameters for the different candidate distributions as a function of threshold.

**Figure 9.**
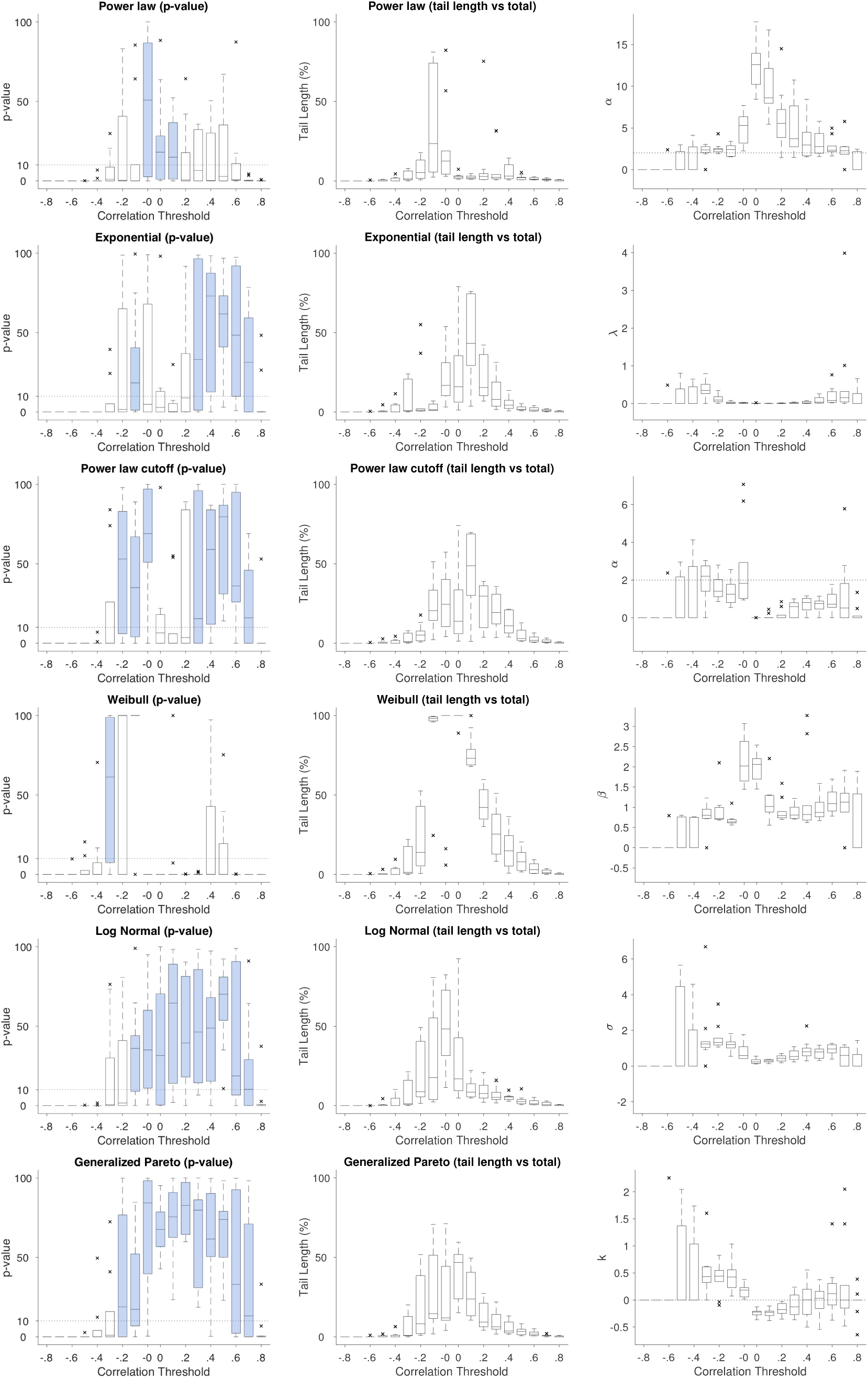
10K data set. Left: population averaged goodness-of-fit tests. Center: percentage of the tail of the distribution explained by the model (center) across different thresholds for each of the six distributions. Horizontal dashed lines in the box-plots indicate the acceptance criteria for a model to be considered plausible (p-value*>* 10).The central mark is the median, the edges of the boxes are the 25*^th^* and 75*^th^* percentiles. Cross marks correspond to outliers. Right: Estimated scaling parameters for the different candidate distributions as a function of threshold.

**Figure 10.**
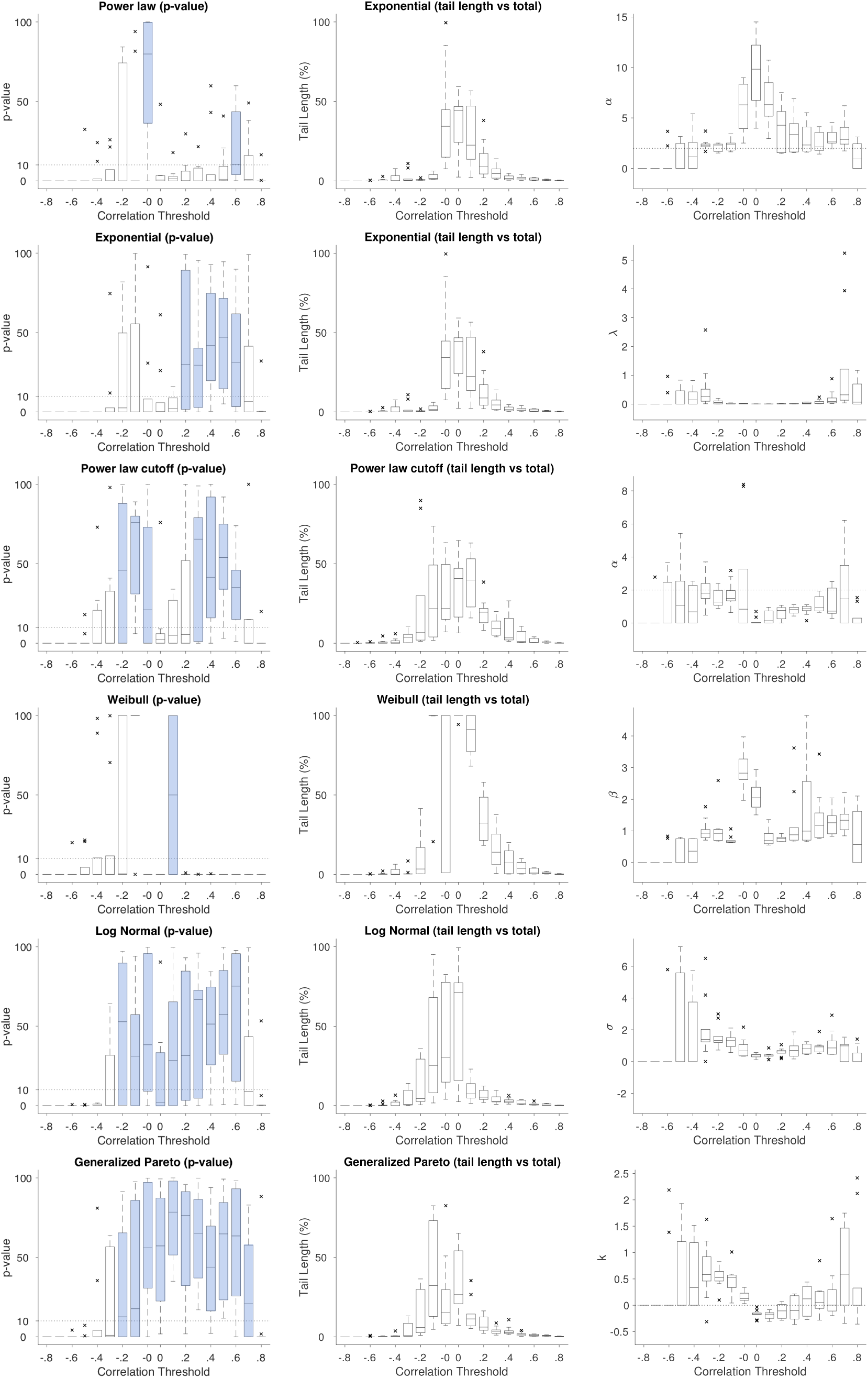
20K data set. Left: population averaged goodness-of-fit tests. Center: percentage of the tail of the distribution explained by the model (center) across different thresholds for each of the six distributions. Horizontal dashed lines in the box-plots indicate the acceptance criteria for a model to be considered plausible (p-value*>* 10).The central mark is the median, the edges of the boxes are the 25*^th^* and 75*^th^* percentiles. Cross marks correspond to outliers. Right: Estimated scaling parameters for the different candidate distributions as a function of threshold

**Figure 11.**
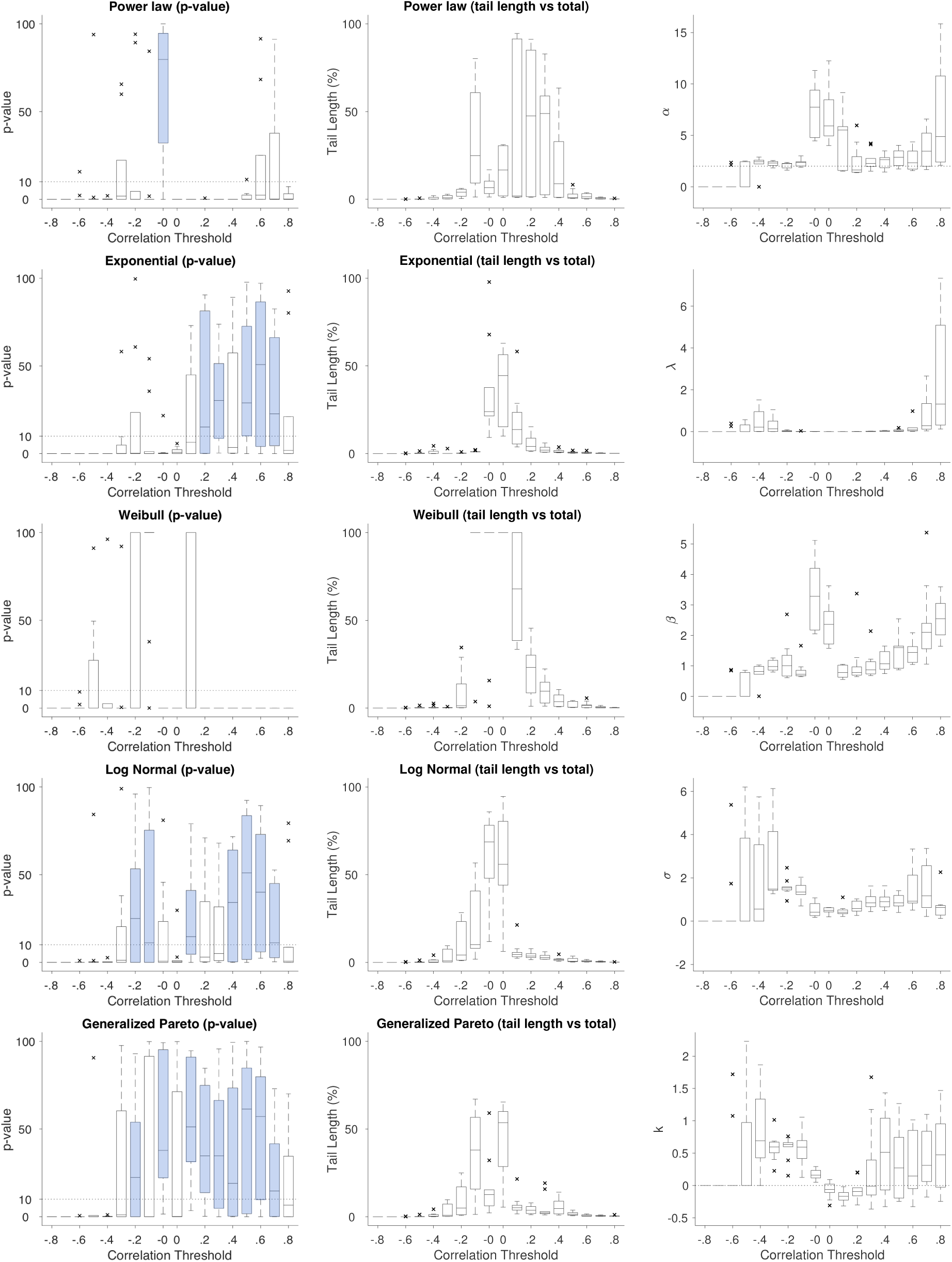
50K data set. Left: population averaged goodness-of-fit tests. Center: percentage of the tail of the distribution explained by the model (center) across different thresholds for each of the five distributions. Horizontal dashed lines in the box-plots indicate the acceptance criteria for a model to be considered plausible (p-value*>* 10).The central mark is the median, the edges of the boxes are the 25*^th^* and 75*^th^* percentiles. Cross marks correspond to outliers. Right: Estimated scaling parameters for the different candidate distributions as a function of threshold.

**Figure 12.**
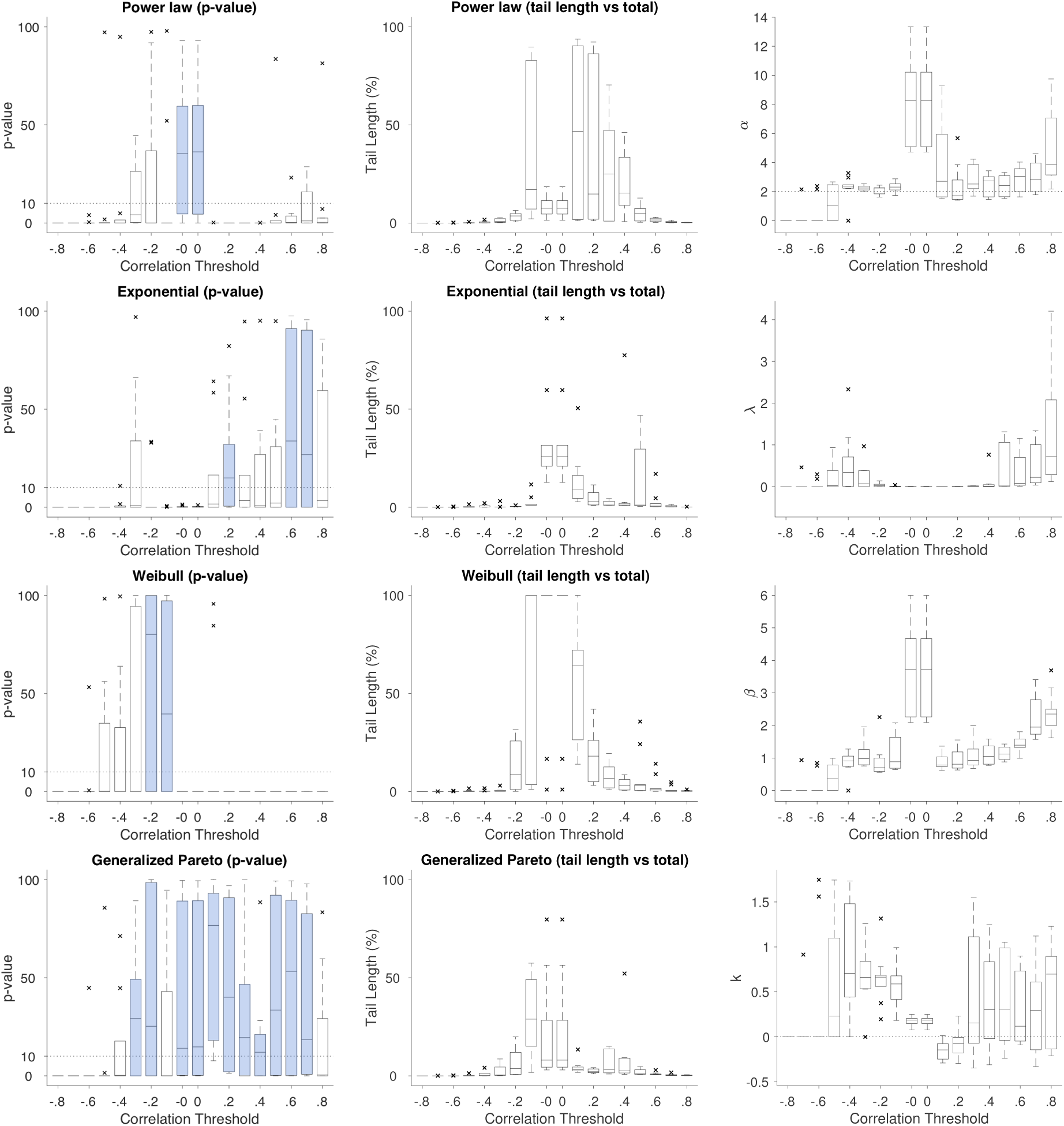
80K data set. Left: population averaged goodness-of-fit tests. Center: percentage of the tail of the distribution explained by the model (center) across different thresholds for each of the four distributions. Horizontal dashed lines in the box-plots indicate the acceptance criteria for a model to be considered plausible (p-value*>* 10).The central mark is the median, the edges of the boxes are the 25*^th^* and 75*^th^* percentiles. Cross marks correspond to outliers. Right: Estimated scaling parameters for the different candidate distributions as a function of threshold.

**Table 5.** Fit results of the exponential distribution for the 1K resolution dataset.

| Thr | $\lambda$ | $x_{min}$ | TL | KS | p-value | TL<sub>r</sub> |
| --- | --- | --- | --- | --- | --- | --- |
| +0.8 | 0.068 | 0.404 | 25.500 | 0.029 | 0.000 | 0.747 |
| +0.7 | 0.065 | 1.093 | 54.500 | 0.063 | 0.009 | 0.695 |
| +0.6 | 0.110 | 2.221 | 161.500 | 0.063 | 0.176 | 0.650 |
| +0.5 | 0.060 | 2.180 | 187.000 | 0.061 | 0.068 | 0.667 |
| +0.4 | 0.053 | 24.811 | 255.000 | 0.061 | 0.205 | 0.541 |
| +0.3 | 0.047 | 43.658 | 356.500 | 0.071 | 0.035 | 0.490 |
| +0.2 | 0.041 | 103.496 | 163.000 | 0.069 | 0.122 | 0.199 |
| +0.1 | 0.063 | 172.515 | 113.500 | 0.071 | 0.444 | 0.116 |
| +0.0 | 0.108 | 5.186 | 234.500 | 0.062 | 0.508 | 0.233 |
| -0.0 | 0.108 | 5.186 | 234.500 | 0.062 | 0.520 | 0.233 |
| -0.1 | 0.224 | 1.395 | 267.500 | 0.116 | 0.042 | 0.383 |
| -0.2 | 0.555 | 0.383 | 71.500 | 0.202 | 0.000 | 0.396 |
| -0.3 | 0.000 | 0.000 | 0.000 | 0.000 | 0.000 | 0.898 |
| -0.4 | 0.000 | 0.000 | 0.000 | 0.000 | 0.000 | - |
| -0.5 | 0.000 | 0.000 | 0.000 | 0.000 | 0.000 | - |
| -0.6 | 0.000 | 0.000 | 0.000 | 0.000 | 0.000 | - |

**Table 6.**
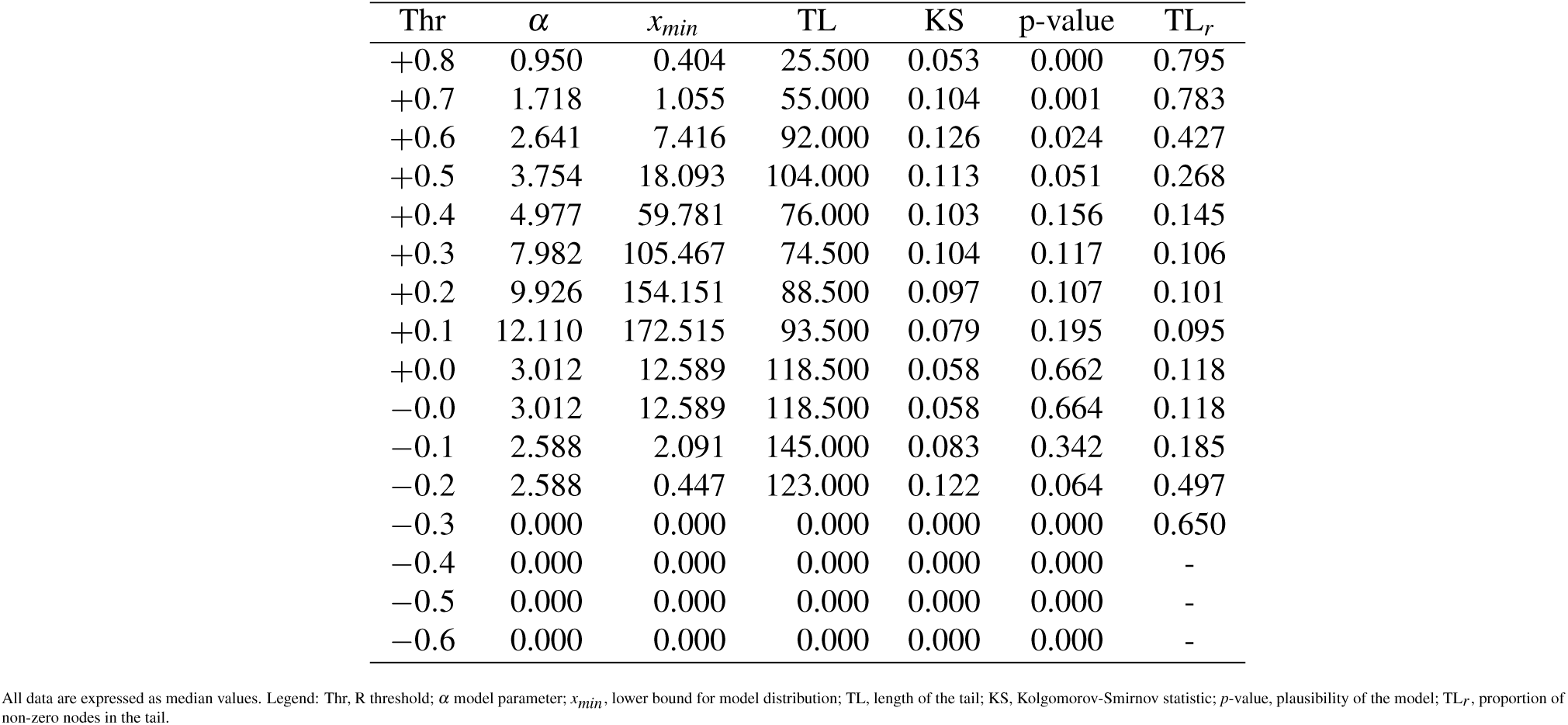
Fit results of the power law distribution for the 1K resolution dataset.

**Table 7.** Fit results of the generalized Pareto distribution for the 1K resolution dataset.

| Thr | $k$ | $\sigma$ | $x_{min}$ | TL | KS | p-value | TL<sub>r</sub> |
| --- | --- | --- | --- | --- | --- | --- | --- |
| +0.8 | 0.000 | 0.696 | 0.403 | 25.500 | 0.029 | 0.000 | 0.831 |
| +0.7 | -0.072 | 5.662 | 1.847 | 53.500 | 0.051 | 0.122 | 0.568 |
| +0.6 | -0.230 | 12.132 | 7.004 | 100.500 | 0.058 | 0.722 | 0.493 |
| +0.5 | -0.211 | 22.787 | 9.726 | 147.500 | 0.035 | 0.868 | 0.404 |
| +0.4 | -0.340 | 30.337 | 13.315 | 229.500 | 0.031 | 0.889 | 0.453 |
| +0.3 | -0.440 | 36.590 | 53.520 | 211.500 | 0.030 | 0.910 | 0.314 |
| +0.2 | -0.399 | 39.570 | 84.198 | 226.500 | 0.031 | 0.830 | 0.274 |
| +0.1 | -0.420 | 51.221 | 107.558 | 354.000 | 0.033 | 0.784 | 0.357 |
| +0.0 | 0.405 | 3.788 | 2.983 | 540.500 | 0.022 | 0.615 | 0.539 |
| -0.0 | 0.405 | 3.788 | 2.983 | 540.500 | 0.022 | 0.610 | 0.539 |
| -0.1 | 0.469 | 2.028 | 1.061 | 344.000 | 0.036 | 0.599 | 0.463 |
| -0.2 | 0.521 | 0.827 | 0.551 | 88.500 | 0.070 | 0.292 | 0.401 |
| -0.3 | 0.000 | 0.000 | 0.000 | 0.000 | 0.000 | 0.000 | 0.686 |
| -0.4 | 0.000 | 0.000 | 0.000 | 0.000 | 0.000 | 0.000 | - |
| -0.5 | 0.000 | 0.000 | 0.000 | 0.000 | 0.000 | 0.000 | - |
| -0.6 | 0.000 | 0.000 | 0.000 | 0.000 | 0.000 | 0.000 | - |

**Table 8.** Fit results of the log-normal distribution for the 1K resolution dataset.

| Thr | $\mu$ | $\sigma$ | $x_{min}$ | TL | KS | p-value | TL<sub>r</sub> |
| --- | --- | --- | --- | --- | --- | --- | --- |
| +0.8 | 0.000 | 0.383 | 0.404 | 25.500 | 0.028 | 0.000 | 0.786 |
| +0.7 | 1.025 | 0.466 | 0.718 | 53.000 | 0.051 | 0.002 | 0.465 |
| +0.6 | 1.919 | 0.678 | 1.928 | 138.000 | 0.058 | 0.228 | 0.599 |
| +0.5 | 3.189 | 0.476 | 14.200 | 157.500 | 0.043 | 0.642 | 0.394 |
| +0.4 | 4.112 | 0.281 | 42.950 | 149.500 | 0.037 | 0.903 | 0.279 |
| +0.3 | 4.603 | 0.200 | 79.863 | 145.000 | 0.036 | 0.695 | 0.233 |
| +0.2 | 4.981 | 0.162 | 124.291 | 152.000 | 0.036 | 0.862 | 0.196 |
| +0.1 | 5.061 | 0.145 | 132.611 | 192.000 | 0.036 | 0.835 | 0.194 |
| +0.0 | 1.746 | 1.153 | 4.474 | 540.500 | 0.028 | 0.390 | 0.539 |
| -0.0 | 1.746 | 1.153 | 4.474 | 540.500 | 0.028 | 0.392 | 0.539 |
| -0.1 | 0.132 | 1.190 | 0.896 | 364.500 | 0.035 | 0.782 | 0.490 |
| -0.2 | -0.238 | 1.072 | 0.567 | 69.000 | 0.059 | 0.307 | 0.382 |
| -0.3 | 0.000 | 0.000 | 0.000 | 0.000 | 0.000 | 0.000 | 0.823 |
| -0.4 | 0.000 | 0.000 | 0.000 | 0.000 | 0.000 | 0.000 | - |
| -0.5 | 0.000 | 0.000 | 0.000 | 0.000 | 0.000 | 0.000 | - |
| -0.6 | 0.000 | 0.000 | 0.000 | 0.000 | 0.000 | 0.000 | - |

**Table 9.** Fit results of the Weibull distribution for the 1K resolution dataset.

| Thr | $b$ | $a$ | $x_{min}$ | TL | KS | p-value | TL<sub>r</sub> |
| --- | --- | --- | --- | --- | --- | --- | --- |
| +0.8 | 0.595 | 1.194 | 0.404 | 25.500 | 0.038 | 0.000 | 0.775 |
| +0.7 | 1.175 | 6.292 | 0.723 | 61.500 | 0.062 | 0.000 | 0.843 |
| +0.6 | 1.358 | 16.338 | 0.951 | 166.500 | 0.061 | 0.000 | 0.862 |
| +0.5 | 1.337 | 25.943 | 2.938 | 326.000 | 0.051 | 0.000 | 0.857 |
| +0.4 | 1.332 | 48.639 | 3.155 | 484.500 | 0.046 | 0.000 | 0.898 |
| +0.3 | 2.173 | 83.217 | 21.348 | 527.000 | 0.041 | 0.000 | 0.731 |
| +0.2 | 2.581 | 110.484 | 24.139 | 639.500 | 0.043 | 0.000 | 0.743 |
| +0.2 | 3.026 | 145.226 | 44.496 | 736.000 | 0.040 | 0.000 | 0.745 |
| +0.0 | 1.385 | 10.267 | 1.406 | 1002.000 | 0.109 | 0.000 | 1.000 |
| -0.0 | 1.385 | 10.267 | 1.406 | 1002.000 | 0.109 | 0.000 | 1.000 |
| -0.1 | 0.783 | 2.090 | 0.120 | 693.000 | 0.120 | 0.895 | 0.883 |
| -0.2 | 0.942 | 1.505 | 0.249 | 125.000 | 0.184 | 0.000 | 0.659 |
| -0.3 | 0.000 | 0.000 | 0.000 | 0.000 | 0.000 | 0.000 | 0.730 |
| -0.4 | 0.000 | 0.000 | 0.000 | 0.000 | 0.000 | 0.000 | - |
| -0.5 | 0.000 | 0.000 | 0.000 | 0.000 | 0.000 | 0.000 | - |
| -0.6 | 0.000 | 0.000 | 0.000 | 0.000 | 0.000 | 0.000 | - |

**Table 10.** Fit results of the power law with exponential cutoff distribution for the 1K resolution dataset.

| Thr | $\alpha$ | $\lambda$ | $x_{min}$ | TL | KS | p-value | TL<sub>r</sub> |
| --- | --- | --- | --- | --- | --- | --- | --- |
| +0.8 | 0.000 | 0.049 | 0.404 | 25.500 | 0.030 | 0.000 | 0.864 |
| +0.7 | 0.076 | 0.051 | 0.715 | 57.500 | 0.060 | 0.000 | 0.841 |
| +0.6 | 0.398 | 0.072 | 1.602 | 241.500 | 0.060 | 0.105 | 0.805 |
| +0.5 | 0.141 | 0.047 | 0.819 | 288.500 | 0.051 | 0.390 | 0.867 |
| +0.4 | 0.006 | 0.026 | 0.484 | 525.000 | 0.068 | 0.225 | 0.938 |
| +0.3 | 0.000 | 0.016 | 0.343 | 704.500 | 0.101 | 0.270 | 0.975 |
| +0.2 | 0.000 | 0.013 | 0.360 | 836.500 | 0.140 | 0.330 | 0.965 |
| +0.1 | 0.000 | 0.010 | 0.124 | 977.500 | 0.159 | 0.350 | 0.988 |
| +0.0 | 1.048 | 0.044 | 3.174 | 645.000 | 0.025 | 0.605 | 0.646 |
| -0.0 | 1.048 | 0.044 | 3.174 | 645.000 | 0.025 | 0.615 | 0.646 |
| -0.1 | 1.548 | 0.051 | 0.834 | 399.000 | 0.041 | 0.485 | 0.475 |
| -0.2 | 1.883 | 0.060 | 0.639 | 108.500 | 0.077 | 0.335 | 0.372 |
| -0.3 | 0.000 | 0.000 | 0.000 | 0.000 | 0.000 | 0.000 | 0.500 |
| -0.4 | 0.000 | 0.000 | 0.000 | 0.000 | 0.000 | 0.000 | - |
| -0.5 | 0.000 | 0.000 | 0.000 | 0.000 | 0.000 | 0.000 | - |
| -0.6 | 0.000 | 0.000 | 0.000 | 0.000 | 0.000 | 0.000 | - |

**Table 11.** Fit results of the exponential distribution for the 5K resolution dataset.

| Thr | $\lambda$ | $x_{min}$ | TL | KS | p-value | TL<sub>r</sub> |
| --- | --- | --- | --- | --- | --- | --- |
| +0.8 | 0.000 | 0.000 | 0.000 | 0.000 | 0.000 | 0.665 |
| +0.7 | 0.055 | 1.877 | 56.000 | 0.042 | 0.254 | 0.436 |
| +0.6 | 0.085 | 4.515 | 134.500 | 0.078 | 0.194 | 0.403 |
| +0.5 | 0.042 | 10.185 | 303.000 | 0.046 | 0.533 | 0.374 |
| +0.4 | 0.023 | 12.670 | 597.000 | 0.043 | 0.167 | 0.392 |
| +0.3 | 0.016 | 26.930 | 455.500 | 0.034 | 0.414 | 0.229 |
| +0.2 | 0.014 | 56.294 | 949.000 | 0.032 | 0.116 | 0.258 |
| +0.1 | 0.010 | 103.691 | 1866.500 | 0.047 | 0.001 | 0.374 |
| +0.0 | 0.038 | 58.758 | 1190.500 | 0.023 | 0.429 | 0.238 |
| -0.0 | 0.038 | 58.758 | 1190.500 | 0.023 | 0.425 | 0.238 |
| -0.1 | 0.056 | 14.940 | 368.500 | 0.048 | 0.233 | 0.080 |
| -0.2 | 0.206 | 1.418 | 227.500 | 0.137 | 0.000 | 0.274 |
| -0.3 | 0.556 | 0.317 | 106.000 | 0.235 | 0.000 | 0.789 |
| -0.4 | 0.000 | 0.000 | 0.000 | 0.000 | 0.000 | 0.754 |
| -0.5 | 0.000 | 0.000 | 0.000 | 0.000 | 0.000 | 0.892 |
| -0.6 | 0.000 | 0.000 | 0.000 | 0.000 | 0.000 | - |

All data are expressed as median values. Legend: Thr, R threshold; $\lambda$ model parameter; $x_{min}$ , lower bound for model distribution; TL, length of the tail; KS, Kolgomorov-Smirnov statistic; $p$ -value, plausibility of the model; TL<sub>r</sub>, proportion of non-zero nodes in the tail.

**Table 12.** Fit results of the power law distribution for the 5K resolution dataset.

| Thr | $\alpha$ | $x_{min}$ | TL | KS | p-value | TL<sub>r</sub> |
| --- | --- | --- | --- | --- | --- | --- |
| +0.8 | 0.000 | 0.000 | 0.000 | 0.000 | 0.000 | 0.760 |
| +0.7 | 1.705 | 1.414 | 59.000 | 0.084 | 0.001 | 0.310 |
| +0.6 | 2.511 | 2.750 | 81.000 | 0.115 | 0.038 | 0.370 |
| +0.5 | 2.819 | 19.770 | 132.500 | 0.094 | 0.025 | 0.192 |
| +0.4 | 3.783 | 93.027 | 141.500 | 0.086 | 0.087 | 0.109 |
| +0.3 | 6.883 | 170.489 | 113.500 | 0.077 | 0.270 | 0.042 |
| +0.2 | 9.258 | 284.261 | 125.000 | 0.077 | 0.229 | 0.031 |
| +0.1 | 12.458 | 517.130 | 155.000 | 0.074 | 0.108 | 0.031 |
| +0.0 | 4.907 | 76.779 | 384.500 | 0.033 | 0.599 | 0.077 |
| -0.0 | 4.907 | 76.779 | 384.500 | 0.033 | 0.607 | 0.077 |
| -0.1 | 2.537 | 9.146 | 592.500 | 0.065 | 0.114 | 0.125 |
| -0.2 | 2.558 | 2.109 | 611.500 | 0.069 | 0.004 | 0.353 |
| -0.3 | 2.411 | 0.802 | 83.000 | 0.112 | 0.001 | 0.407 |
| -0.4 | 0.000 | 0.000 | 0.000 | 0.000 | 0.000 | 0.473 |
| -0.5 | 0.000 | 0.000 | 0.000 | 0.000 | 0.000 | 0.938 |
| -0.6 | 0.000 | 0.000 | 0.000 | 0.000 | 0.000 | - |

All data are expressed as median values. Legend: Thr, R threshold; $\alpha$ model parameter; $x_{min}$ , lower bound for model distribution; TL, length of the tail; KS, Kolgomorov-Smirnov statistic; $p$ -value, plausibility of the model; TL<sub>r</sub>, proportion of non-zero nodes in the tail.

**Table 13.** Fit results of the generalized Pareto distribution for the 5K resolution dataset.

| Thr | $k$ | $\sigma$ | $x_{min}$ | TL | KS | p-value | TL<sub>r</sub> |
| --- | --- | --- | --- | --- | --- | --- | --- |
| +0.8 | 0.000 | 0.000 | 0.000 | 0.000 | 0.000 | 0.000 | 0.635 |
| +0.7 | 0.000 | 4.044 | 1.485 | 68.500 | 0.037 | 0.102 | 0.412 |
| +0.6 | 0.038 | 11.339 | 4.468 | 123.000 | 0.049 | 0.414 | 0.408 |
| +0.5 | 0.011 | 23.610 | 6.611 | 341.500 | 0.031 | 0.649 | 0.390 |
| +0.4 | -0.146 | 59.947 | 34.465 | 497.500 | 0.026 | 0.671 | 0.350 |
| +0.3 | -0.231 | 93.821 | 45.148 | 588.000 | 0.025 | 0.573 | 0.231 |
| +0.2 | -0.251 | 140.834 | 50.755 | 1017.500 | 0.016 | 0.618 | 0.296 |
| +0.1 | -0.307 | 144.573 | 102.543 | 1583.000 | 0.014 | 0.653 | 0.316 |
| +0.0 | 0.081 | 21.723 | 45.926 | 1561.500 | 0.014 | 0.805 | 0.312 |
| -0.0 | 0.081 | 21.723 | 45.926 | 1561.500 | 0.014 | 0.787 | 0.312 |
| -0.1 | 0.196 | 11.532 | 9.101 | 643.500 | 0.018 | 0.883 | 0.133 |
| -0.2 | 0.376 | 1.841 | 0.894 | 355.000 | 0.036 | 0.568 | 0.430 |
| -0.3 | 0.664 | 0.586 | 0.942 | 67.500 | 0.098 | 0.005 | 0.399 |
| -0.4 | 0.000 | 0.000 | 0.000 | 0.000 | 0.000 | 0.000 | 0.280 |
| -0.5 | 0.000 | 0.000 | 0.000 | 0.000 | 0.000 | 0.000 | 0.825 |
| -0.6 | 0.000 | 0.000 | 0.000 | 0.000 | 0.000 | 0.000 | - |

All data are expressed as median values. Legend: Thr, R threshold; $k$ , $\sigma$ model parameters; $x_{min}$ , lower bound for model distribution; TL, length of the tail; KS, Kolgomorov-Smirnov statistic; $p$ -value, plausibility of the model; TL<sub>r</sub>, proportion of non-zero nodes in the tail.

**Table 14.** Fit results of the log-normal distribution for the 5K resolution dataset.

| Thr | $\mu$ | $\sigma$ | $x_{min}$ | TL | KS | p-value | TL<sub>r</sub> |
| --- | --- | --- | --- | --- | --- | --- | --- |
| +0.8 | 0.000 | 0.000 | 0.000 | 0.000 | 0.000 | 0.000 | 0.581 |
| +0.7 | 1.154 | 0.484 | 1.484 | 57.000 | 0.043 | 0.178 | 0.396 |
| +0.6 | 1.840 | 0.940 | 1.967 | 161.500 | 0.043 | 0.268 | 0.430 |
| +0.5 | 3.065 | 0.762 | 7.883 | 228.500 | 0.032 | 0.842 | 0.289 |
| +0.4 | 3.501 | 0.575 | 27.431 | 547.000 | 0.031 | 0.462 | 0.273 |
| +0.3 | 4.234 | 0.539 | 47.040 | 617.000 | 0.026 | 0.477 | 0.238 |
| +0.2 | 5.526 | 0.302 | 197.403 | 641.000 | 0.024 | 0.628 | 0.172 |
| +0.1 | 5.854 | 0.226 | 258.005 | 442.500 | 0.021 | 0.660 | 0.089 |
| +0.0 | 3.774 | 0.530 | 29.803 | 2801.000 | 0.011 | 0.838 | 0.556 |
| -0.0 | 3.774 | 0.530 | 29.803 | 2801.000 | 0.011 | 0.808 | 0.556 |
| -0.1 | 2.074 | 0.948 | 6.704 | 634.500 | 0.026 | 0.710 | 0.139 |
| -0.2 | 0.469 | 1.080 | 0.934 | 615.000 | 0.041 | 0.298 | 0.487 |
| -0.3 | -1.652 | 1.487 | 0.350 | 115.000 | 0.123 | 0.002 | 0.657 |
| -0.4 | 0.000 | 0.000 | 0.000 | 0.000 | 0.000 | 0.000 | 0.518 |
| -0.5 | 0.000 | 0.000 | 0.000 | 0.000 | 0.000 | 0.000 | -0.010 |
| -0.6 | 0.000 | 0.000 | 0.000 | 0.000 | 0.000 | 0.000 | - |

All data are expressed as median values. Legend: Thr, R threshold; $\mu$ , $\sigma$ model parameters; $x_{min}$ , lower bound for model distribution; TL, length of the tail; KS, Kolgomorov-Smirnov statistic; $p$ -value, plausibility of the model; TL<sub>r</sub>, proportion of non-zero nodes in the tail.

**Table 15.** Fit results of the Weibull distribution for the 5K resolution dataset.

| Thr | $b$ | $a$ | $x_{min}$ | TL | KS | p-value | TL<sub>r</sub> |
| --- | --- | --- | --- | --- | --- | --- | --- |
| +0.8 | 0.000 | 0.000 | 0.000 | 0.000 | 0.000 | 0.000 | 0.773 |
| +0.7 | 0.926 | 4.464 | 0.712 | 85.000 | 0.068 | 0.000 | 0.788 |
| +0.6 | 1.085 | 8.981 | 0.646 | 244.000 | 0.084 | 0.000 | 0.693 |
| +0.5 | 0.926 | 16.819 | 0.563 | 554.000 | 0.074 | 0.000 | 0.769 |
| +0.4 | 0.835 | 34.284 | 0.656 | 1071.000 | 0.064 | 0.026 | 0.823 |
| +0.3 | 0.818 | 50.100 | 0.782 | 1746.000 | 0.052 | 0.000 | 0.763 |
| +0.2 | 1.062 | 117.434 | 5.490 | 2610.000 | 0.043 | 0.000 | 0.698 |
| +0.1 | 1.526 | 216.007 | 21.726 | 3822.000 | 0.038 | 0.000 | 0.763 |
| +0.0 | 1.974 | 56.210 | 14.462 | 5009.500 | 0.107 | 0.000 | 1.000 |
| -0.0 | 1.974 | 56.210 | 14.462 | 5009.500 | 0.107 | 0.000 | 1.000 |
| -0.1 | 0.700 | 5.459 | 0.110 | 4193.000 | 0.094 | 1.000 | 0.915 |
| -0.2 | 0.782 | 2.817 | 0.230 | 680.500 | 0.141 | 0.997 | 0.781 |
| -0.3 | 0.824 | 1.666 | 0.338 | 104.500 | 0.203 | 0.004 | 0.718 |
| -0.4 | 0.000 | 0.000 | 0.000 | 0.000 | 0.000 | 0.000 | 0.550 |
| -0.5 | 0.000 | 0.000 | 0.000 | 0.000 | 0.000 | 0.000 | 0.492 |
| -0.6 | 0.000 | 0.000 | 0.000 | 0.000 | 0.000 | 0.000 | - |

All data are expressed as median values. Legend: Thr, R threshold; $b$ , $a$ model parameters; $x_{min}$ , lower bound for model distribution; TL, length of the tail; KS, Kolgomorov-Smirnov statistic; $p$ -value, plausibility of the model; TL<sub>r</sub>, proportion of non-zero nodes in the tail.

**Table 16.** Fit results of the power law with exponential cutoff distribution for the 5K resolution dataset.

| Thr | $\alpha$ | $\lambda$ | $x_{min}$ | TL | KS | p-value | TL<sub>r</sub> |
| --- | --- | --- | --- | --- | --- | --- | --- |
| +0.8 | 0.000 | 0.000 | 0.000 | 0.000 | 0.000 | 0.000 | 0.673 |
| +0.7 | 0.100 | 0.046 | 1.122 | 60.500 | 0.040 | 0.155 | 0.466 |
| +0.6 | 0.583 | 0.065 | 1.883 | 193.000 | 0.056 | 0.300 | 0.621 |
| +0.5 | 0.484 | 0.037 | 3.328 | 541.500 | 0.030 | 0.590 | 0.551 |
| +0.4 | 0.592 | 0.016 | 1.898 | 1117.000 | 0.022 | 0.390 | 0.671 |
| +0.3 | 0.314 | 0.010 | 3.231 | 1833.500 | 0.024 | 0.120 | 0.704 |
| +0.2 | 0.188 | 0.007 | 3.829 | 2778.000 | 0.035 | 0.015 | 0.722 |
| +0.1 | 0.016 | 0.006 | 3.789 | 4263.500 | 0.067 | 0.015 | 0.853 |
| +0.0 | 0.748 | 0.026 | 23.766 | 3764.500 | 0.041 | 0.185 | 0.748 |
| -0.0 | 0.748 | 0.026 | 23.766 | 3764.500 | 0.041 | 0.185 | 0.748 |
| -0.1 | 1.475 | 0.026 | 4.251 | 1593.000 | 0.019 | 0.470 | 0.340 |
| -0.2 | 1.837 | 0.029 | 1.810 | 517.000 | 0.043 | 0.070 | 0.272 |
| -0.3 | 2.022 | 0.000 | 0.836 | 80.000 | 0.098 | 0.000 | 0.353 |
| -0.4 | 0.000 | 0.000 | 0.000 | 0.000 | 0.000 | 0.000 | 0.473 |
| -0.5 | 0.000 | 0.000 | 0.000 | 0.000 | 0.000 | 0.000 | 0.946 |
| -0.6 | 0.000 | 0.000 | 0.000 | 0.000 | 0.000 | 0.000 | - |

All data are expressed as median values. Legend: Thr, R threshold; $b$ , $a$ model parameters; $x_{min}$ , lower bound for model distribution; TL, length of the tail; KS, Kolgomorov-Smirnov statistic; $p$ -value, plausibility of the model; TL<sub>r</sub>, proportion of non-zero nodes in the tail.

**Table 17.**
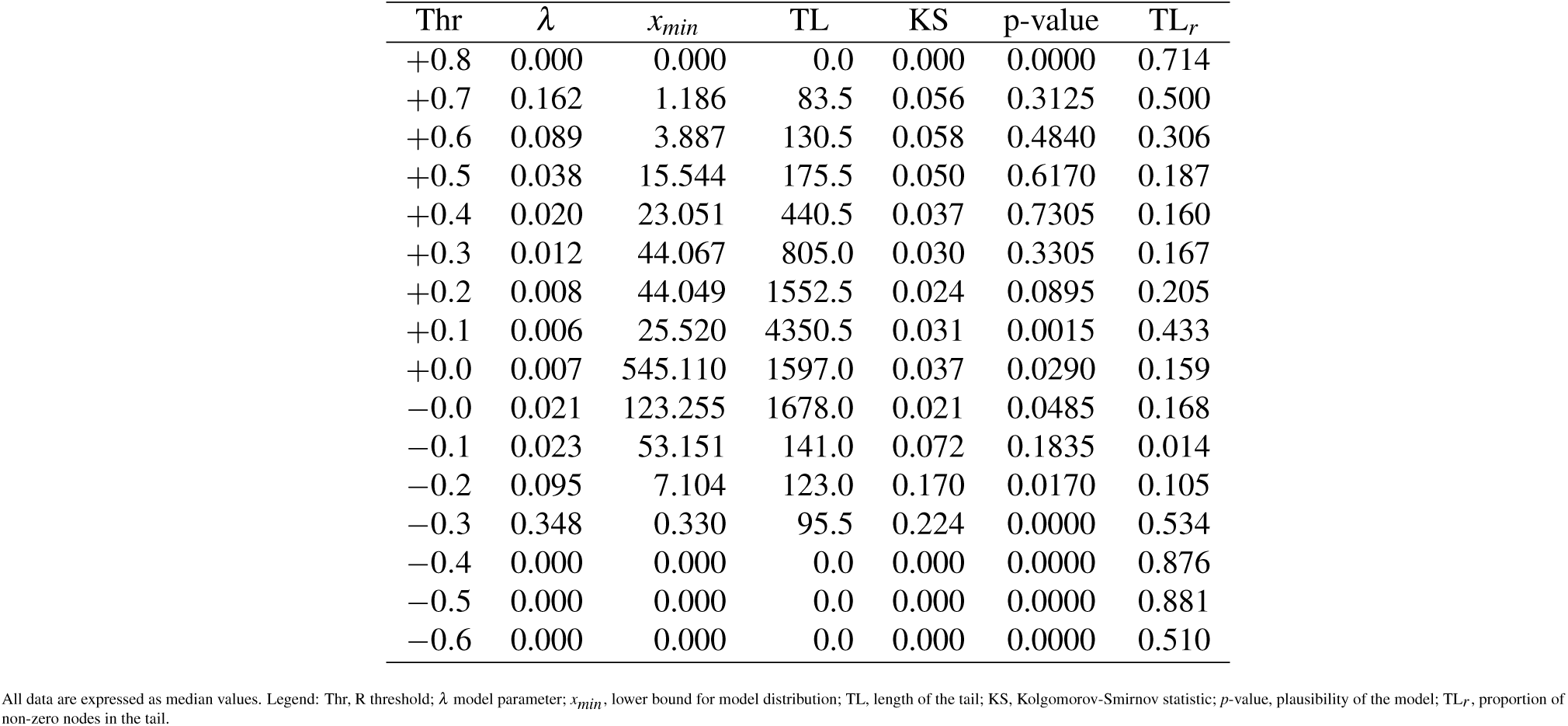
Fit results of the exponential distribution for the 10K resolution dataset.

**Table 18.**
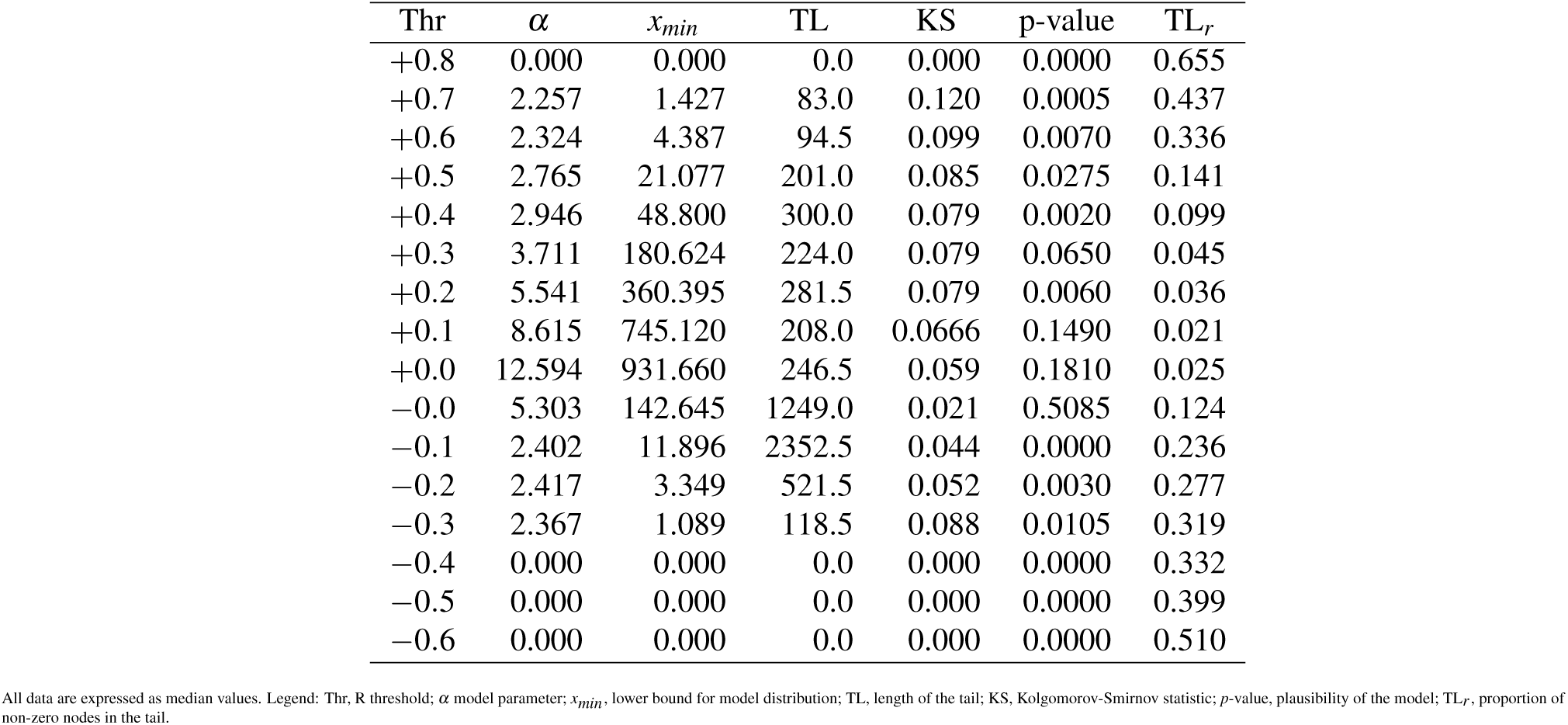
Fit results of the power law distribution for the 10K resolution dataset.

**Table 19.** Fit results of the generalized Pareto distribution for the 10K resolution dataset.

| Thr | $k$ | $\sigma$ | $x_{min}$ | TL | KS | p-value | TL<sub>r</sub> |
| --- | --- | --- | --- | --- | --- | --- | --- |
| +0.8 | 0.000 | 0.000 | 0.000 | 0.0 | 0.000 | 0.0000 | 0.696 |
| +0.7 | 0.000 | 3.663 | 1.539 | 58.5 | 0.049 | 0.1315 | 0.531 |
| +0.6 | 0.118 | 11.094 | 2.943 | 172.5 | 0.048 | 0.3305 | 0.440 |
| +0.5 | 0.031 | 28.656 | 20.072 | 338.5 | 0.034 | 0.7380 | 0.229 |
| +0.4 | 0.001 | 62.035 | 34.191 | 372.5 | 0.027 | 0.6155 | 0.174 |
| +0.3 | -0.128 | 102.619 | 85.215 | 638.5 | 0.021 | 0.7975 | 0.176 |
| +0.2 | -0.178 | 152.365 | 136.945 | 938.5 | 0.014 | 0.8275 | 0.135 |
| +0.1 | -0.226 | 241.640 | 201.475 | 2387.0 | 0.010 | 0.7555 | 0.238 |
| +0.0 | -0.224 | 218.500 | 380.050 | 4681.5 | 0.009 | 0.6750 | 0.467 |
| -0.0 | 0.182 | 32.619 | 127.480 | 1195.5 | 0.012 | 0.8425 | 0.119 |
| -0.1 | 0.426 | 12.883 | 9.690 | 1469.0 | 0.020 | 0.1725 | 0.149 |
| -0.2 | 0.439 | 2.549 | 0.763 | 820.5 | 0.030 | 0.1890 | 0.457 |
| -0.3 | 0.433 | 0.996 | 0.649 | 101.5 | 0.062 | 0.0105 | 0.405 |
| -0.4 | 0.000 | 0.000 | 0.000 | 0.0 | 0.000 | 0.0000 | 0.253 |
| -0.5 | 0.000 | 0.000 | 0.000 | 0.0 | 0.000 | 0.0000 | 1.000 |
| -0.6 | 0.000 | 0.000 | 0.000 | 0.0 | 0.000 | 0.0000 | 1.000 |

All data are expressed as median values. Legend: Thr, R threshold; $k$ , $\sigma$ model parameters; $x_{min}$ , lower bound for model distribution; TL, length of the tail; KS, Kolgomorov-Smirnov statistic; $p$ -value, plausibility of the model; TL<sub>r</sub>, proportion of non-zero nodes in the tail.

**Table 20.** Fit results of the log-normal distribution for the 10K resolution dataset.

| Thr | $\mu$ | $\sigma$ | $x_{min}$ | TL | KS | p-value | TL<sub>r</sub> |
| --- | --- | --- | --- | --- | --- | --- | --- |
| +0.8 | 0.000 | 0.000 | 0.000 | 0.000 | 0.000 | 0.000 | 0.639 |
| +0.7 | 0.618 | 0.599 | 1.141 | 58.000 | 0.039 | 0.103 | 0.466 |
| +0.6 | 1.804 | 0.958 | 2.589 | 110.500 | 0.040 | 0.187 | 0.381 |
| +0.5 | 3.363 | 0.794 | 15.168 | 246.500 | 0.033 | 0.702 | 0.223 |
| +0.4 | 3.452 | 0.799 | 24.044 | 543.500 | 0.027 | 0.488 | 0.186 |
| +0.3 | 4.809 | 0.535 | 72.627 | 525.000 | 0.023 | 0.463 | 0.114 |
| +0.2 | 5.754 | 0.434 | 209.015 | 759.500 | 0.023 | 0.395 | 0.100 |
| +0.1 | 6.143 | 0.299 | 418.120 | 852.000 | 0.019 | 0.645 | 0.085 |
| +0.0 | 6.299 | 0.236 | 472.295 | 1684.500 | 0.015 | 0.315 | 0.168 |
| -0.0 | 4.298 | 0.588 | 81.662 | 4837.000 | 0.010 | 0.350 | 0.482 |
| -0.1 | 2.176 | 1.198 | 7.106 | 1764.000 | 0.019 | 0.360 | 0.180 |
| -0.2 | 0.486 | 1.317 | 0.610 | 879.000 | 0.034 | 0.017 | 0.599 |
| -0.3 | -0.243 | 1.237 | 0.543 | 106.000 | 0.072 | 0.005 | 0.592 |
| -0.4 | 0.000 | 0.000 | 0.000 | 0.000 | 0.000 | 0.000 | 0.825 |
| -0.5 | 0.000 | 0.000 | 0.000 | 0.000 | 0.000 | 0.000 | 0.607 |
| -0.6 | 0.000 | 0.000 | 0.000 | 0.000 | 0.000 | 0.000 | - |

All data are expressed as median values. Legend: Thr, R threshold; $\mu$ , $\sigma$ model parameters; $x_{min}$ , lower bound for model distribution; TL, length of the tail; KS, Kolgomorov-Smirnov statistic; $p$ -value, plausibility of the model; TL<sub>r</sub>, proportion of non-zero nodes in the tail.

**Table 21.** Fit results of the Weibull distribution for the 10K resolution dataset.

| Thr | $b$ | $a$ | $x_{min}$ | TL | KS | p-value | $TL_r$ |
| --- | --- | --- | --- | --- | --- | --- | --- |
| +0.8 | 0.000 | 0.000 | 0.000 | 0.000 | 0.000 | 0.000 | 0.732 |
| +0.7 | 1.130 | 4.758 | 0.746 | 100.500 | 0.095 | 0.000 | 0.759 |
| +0.6 | 1.088 | 8.477 | 0.698 | 296.500 | 0.091 | 0.000 | 0.556 |
| +0.5 | 0.874 | 14.914 | 0.654 | 794.000 | 0.087 | 0.000 | 0.611 |
| +0.4 | 0.822 | 41.623 | 1.122 | 1480.000 | 0.082 | 0.001 | 0.608 |
| +0.3 | 0.810 | 57.835 | 1.388 | 2546.500 | 0.071 | 0.000 | 0.536 |
| +0.2 | 0.795 | 74.564 | 1.683 | 4226.000 | 0.055 | 0.000 | 0.555 |
| +0.1 | 1.020 | 202.123 | 7.832 | 7336.000 | 0.045 | 0.000 | 0.731 |
| +0.0 | 2.058 | 425.071 | 40.358 | 10022.500 | 0.032 | 0.000 | 1.000 |
| -0.0 | 2.022 | 111.971 | 35.387 | 10022.500 | 0.133 | 0.000 | 1.000 |
| -0.1 | 0.640 | 4.837 | 0.100 | 9843.500 | 0.102 | 1.000 | 1.000 |
| -0.2 | 0.726 | 3.937 | 0.228 | 1396.500 | 0.131 | 1.000 | 0.781 |
| -0.3 | 0.801 | 2.341 | 0.331 | 140.500 | 0.194 | 0.612 | 0.703 |
| -0.4 | 0.000 | 0.000 | 0.000 | 0.000 | 0.000 | 0.000 | 0.685 |
| -0.5 | 0.000 | 0.000 | 0.000 | 0.000 | 0.000 | 0.000 | 0.619 |
| -0.6 | 0.000 | 0.000 | 0.000 | 0.000 | 0.000 | 0.000 | 0.510 |

All data are expressed as median values. Legend: Thr, R threshold; $b$ , $a$ model parameters; $x_{min}$ , lower bound for model distribution; TL, length of the tail; KS, Kolgomorov-Smirnov statistic; $p$ -value, plausibility of the model; $TL_r$ , proportion of non-zero nodes in the tail.

**Table 22.** Fit results of the power law with exponential cutoff distribution for the 10K resolution dataset.

| Thr | $\alpha$ | $\lambda$ | $x_{min}$ | TL | KS | p-value | $TL_r$ |
| --- | --- | --- | --- | --- | --- | --- | --- |
| +0.8 | 0.000 | 0.000 | 0.000 | 0.000 | 0.000 | 0.000 | 0.714 |
| +0.7 | 0.520 | 0.039 | 1.125 | 77.000 | 0.048 | 0.160 | 0.597 |
| +0.6 | 0.719 | 0.070 | 1.927 | 184.500 | 0.037 | 0.360 | 0.421 |
| +0.5 | 0.747 | 0.028 | 2.208 | 302.000 | 0.029 | 0.795 | 0.358 |
| +0.4 | 0.805 | 0.013 | 1.940 | 1098.500 | 0.019 | 0.590 | 0.421 |
| +0.3 | 0.595 | 0.008 | 4.171 | 1933.000 | 0.017 | 0.155 | 0.429 |
| +0.2 | 0.003 | 0.005 | 40.707 | 2980.500 | 0.018 | 0.035 | 0.374 |
| +0.1 | 0.003 | 0.007 | 35.191 | 4913.000 | 0.029 | 0.000 | 0.488 |
| +0.0 | 0.001 | 0.007 | 541.945 | 1388.000 | 0.034 | 0.065 | 0.138 |
| -0.0 | 1.822 | 0.002 | 98.425 | 2461.000 | 0.012 | 0.690 | 0.245 |
| -0.1 | 1.255 | 0.013 | 9.618 | 2055.500 | 0.016 | 0.350 | 0.207 |
| -0.2 | 1.404 | 0.013 | 1.445 | 520.000 | 0.034 | 0.530 | 0.329 |
| -0.3 | 2.203 | 0.000 | 1.300 | 92.000 | 0.074 | 0.000 | 0.319 |
| -0.4 | 0.000 | 0.000 | 0.000 | 0.000 | 0.000 | 0.000 | 0.332 |
| -0.5 | 0.000 | 0.000 | 0.000 | 0.000 | 0.000 | 0.000 | 0.590 |
| -0.6 | 0.000 | 0.000 | 0.000 | 0.000 | 0.000 | 0.000 | 0.510 |

All data are expressed as median values. Legend: Thr, R threshold; $b$ , $a$ model parameters; $x_{min}$ , lower bound for model distribution; TL, length of the tail; KS, Kolgomorov-Smirnov statistic; $p$ -value, plausibility of the model; $TL_r$ , proportion of non-zero nodes in the tail.

**Table 23.** Fit results of the exponential distribution for the 20K resolution dataset.

| Thr | $\lambda$ | $x_{min}$ | TL | KS | p-value | TL<sub>r</sub> |
| --- | --- | --- | --- | --- | --- | --- |
| +0.8 | 0.067 | 0.402 | 25.500 | 0.043 | 0.000 | 0.537 |
| +0.7 | 0.320 | 1.181 | 109.000 | 0.089 | 0.066 | 0.330 |
| +0.6 | 0.091 | 4.105 | 113.500 | 0.062 | 0.312 | 0.216 |
| +0.5 | 0.037 | 15.185 | 239.500 | 0.048 | 0.471 | 0.101 |
| +0.4 | 0.021 | 54.464 | 327.500 | 0.045 | 0.418 | 0.070 |
| +0.3 | 0.010 | 70.877 | 898.500 | 0.032 | 0.294 | 0.090 |
| +0.2 | 0.006 | 93.748 | 1778.000 | 0.020 | 0.298 | 0.109 |
| +0.1 | 0.004 | 115.855 | 4516.500 | 0.020 | 0.020 | 0.225 |
| +0.0 | 0.004 | 518.180 | 8893.500 | 0.028 | 0.003 | 0.443 |
| −0.0 | 0.014 | 228.770 | 6877.500 | 0.019 | 0.001 | 0.343 |
| −0.1 | 0.021 | 52.362 | 333.000 | 0.093 | 0.000 | 0.017 |
| −0.2 | 0.052 | 14.021 | 92.000 | 0.141 | 0.026 | 0.038 |
| −0.3 | 0.261 | 1.009 | 61.500 | 0.266 | 0.000 | 0.396 |
| −0.4 | 0.144 | 0.200 | 27.500 | 0.142 | 0.000 | 0.739 |
| −0.5 | 0.000 | 0.000 | 0.000 | 0.000 | 0.000 | 0.738 |
| −0.6 | 0.000 | 0.000 | 0.000 | 0.000 | 0.000 | 0.653 |

**Table 24.** Fit results of the power law distribution for the 20K resolution dataset.

| Thr | $\alpha$ | $x_{min}$ | TL | KS | p-value | TL<sub>r</sub> |
| --- | --- | --- | --- | --- | --- | --- |
| +0.8 | 0.927 | 0.402 | 25.500 | 0.056 | 0.000 | 0.568 |
| +0.7 | 2.875 | 2.463 | 71.000 | 0.136 | 0.009 | 0.325 |
| +0.6 | 2.712 | 10.254 | 90.500 | 0.093 | 0.104 | 0.126 |
| +0.5 | 2.133 | 1.637 | 272.000 | 0.073 | 0.010 | 0.199 |
| +0.4 | 2.318 | 46.160 | 833.000 | 0.062 | 0.000 | 0.156 |
| +0.3 | 3.359 | 170.924 | 392.000 | 0.070 | 0.000 | 0.037 |
| +0.2 | 4.275 | 439.870 | 432.500 | 0.062 | 0.001 | 0.025 |
| +0.1 | 6.308 | 908.020 | 467.000 | 0.066 | 0.013 | 0.023 |
| +0.0 | 9.823 | 1170.700 | 492.000 | 0.057 | 0.005 | 0.025 |
| −0.0 | 6.299 | 300.355 | 1437.500 | 0.016 | 0.798 | 0.072 |
| −0.1 | 2.344 | 19.710 | 4518.000 | 0.032 | 0.000 | 0.226 |
| −0.2 | 2.226 | 3.464 | 1502.500 | 0.056 | 0.000 | 0.239 |
| −0.3 | 2.313 | 1.026 | 199.500 | 0.079 | 0.001 | 0.357 |
| −0.4 | 1.141 | 0.200 | 74.500 | 0.020 | 0.000 | 0.315 |
| −0.5 | 0.000 | 0.000 | 0.000 | 0.000 | 0.000 | 0.482 |
| −0.6 | 0.000 | 0.000 | 0.0 | 0.000 | 0.0000 | 0.672 |

All data are expressed as median values. Legend: Thr, R threshold; $\lambda$ , $\alpha$ model parameters; $x_{min}$ , lower bound for model distribution; TL, length of the tail; KS, Kolmogorov-Smirnov statistic; $p$ -value, plausibility of the model; TL<sub>r</sub>, proportion of non-zero nodes in the tail.

**Table 25.** Fit results of generalized Pareto distribution for the 20K resolution dataset.

| Thr | $k$ | $\sigma$ | $x_{min}$ | TL | KS | p-value | $TL_r$ |
| --- | --- | --- | --- | --- | --- | --- | --- |
| +0.8 | 0.000 | 0.023 | 0.400 | 25.500 | 0.027 | 0.000 | 0.448 |
| +0.7 | 0.589 | 1.285 | 1.839 | 136.500 | 0.072 | 0.208 | 0.693 |
| +0.6 | 0.002 | 11.055 | 4.328 | 180.500 | 0.042 | 0.634 | 0.182 |
| +0.5 | 0.051 | 24.567 | 17.856 | 268.000 | 0.036 | 0.647 | 0.103 |
| +0.4 | 0.121 | 60.904 | 29.183 | 519.500 | 0.026 | 0.439 | 0.123 |
| +0.3 | -0.106 | 130.720 | 161.190 | 692.000 | 0.023 | 0.650 | 0.066 |
| +0.2 | -0.109 | 212.785 | 231.480 | 1216.000 | 0.015 | 0.764 | 0.079 |
| +0.1 | -0.171 | 316.070 | 303.185 | 2260.500 | 0.011 | 0.784 | 0.113 |
| +0.0 | -0.167 | 317.320 | 557.405 | 5312.000 | 0.008 | 0.572 | 0.265 |
| -0.0 | 0.125 | 55.521 | 270.430 | 3026.500 | 0.010 | 0.560 | 0.151 |
| -0.1 | 0.527 | 12.322 | 3.984 | 6471.500 | 0.017 | 0.176 | 0.322 |
| -0.2 | 0.525 | 3.316 | 0.785 | 1180.500 | 0.028 | 0.126 | 0.356 |
| -0.3 | 0.580 | 1.452 | 0.880 | 162.000 | 0.056 | 0.010 | 0.491 |
| -0.4 | 0.333 | 0.013 | 0.201 | 45.000 | 0.019 | 0.000 | 0.126 |
| -0.5 | 0.000 | 0.000 | 0.000 | 0.0 | 0.000 | 0.0000 | 0.293 |
| -0.6 | 0.000 | 0.000 | 0.000 | 0.0 | 0.000 | 0.0000 | 0.992 |

All data are expressed as median values. Legend: Thr, R threshold; $k$ , $\sigma$ , $\mu$ model parameters; $x_{min}$ , lower bound for model distribution; TL, length of the tail; KS, Kolgomorov-Smirnov statistic; $p$ -value, plausibility of the model; $TL_r$ , proportion of non-zero nodes in the tail.

**Table 26.**
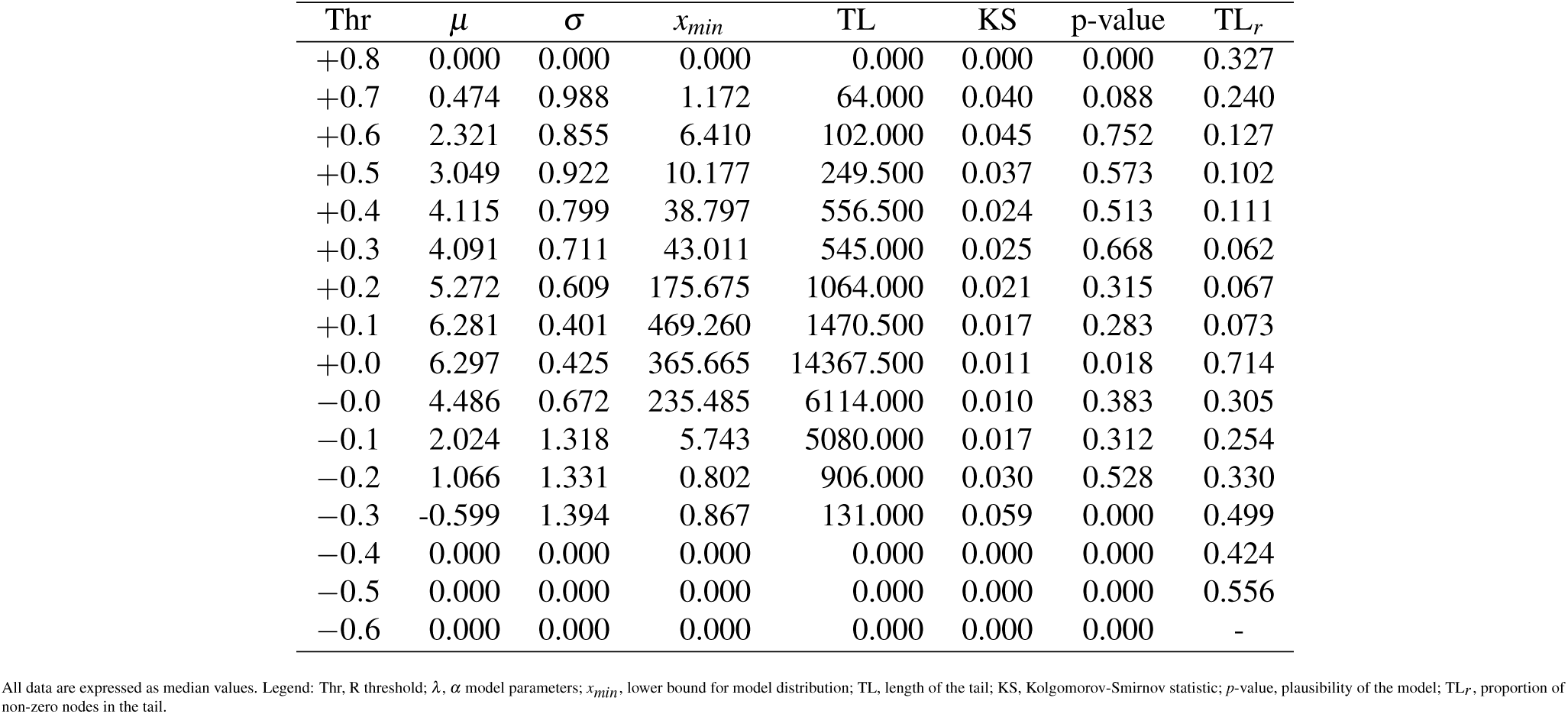
Fit results of the log-normal distribution for the 20K resolution dataset.

**Table 27.** Fit results of the Weibull distribution for the 20K resolution dataset.

| Thr | $\beta$ | $\lambda$ | $x_{min}$ | TL | KS | p-value | TL<sub>r</sub> |
| --- | --- | --- | --- | --- | --- | --- | --- |
| +0.8 | 0.567 | 0.938 | 0.402 | 25.500 | 0.036 | 0.000 | 0.648 |
| +0.7 | 1.335 | 4.353 | 0.803 | 123.000 | 0.133 | 0.000 | 0.500 |
| +0.6 | 1.258 | 13.004 | 1.439 | 183.000 | 0.110 | 0.000 | 0.247 |
| +0.5 | 1.175 | 29.107 | 2.448 | 713.500 | 0.090 | 0.000 | 0.271 |
| +0.4 | 0.993 | 58.500 | 2.273 | 1442.500 | 0.085 | 0.000 | 0.272 |
| +0.3 | 0.878 | 86.394 | 2.574 | 2813.000 | 0.090 | 0.000 | 0.285 |
| +0.2 | 0.768 | 69.888 | 2.397 | 6495.000 | 0.076 | 0.000 | 0.391 |
| +0.1 | 0.697 | 145.778 | 2.101 | 18273.000 | 0.059 | 0.500 | 0.912 |
| +0.0 | 2.049 | 631.567 | 100.867 | 20080.000 | 0.050 | 0.000 | 1.000 |
| -0.0 | 2.824 | 241.419 | 101.626 | 20041.000 | 0.138 | 0.000 | 1.000 |
| -0.1 | 0.657 | 6.909 | 0.100 | 20054.500 | 0.128 | 1.000 | 1.000 |
| -0.2 | 0.925 | 5.265 | 0.687 | 683.000 | 0.131 | 0.004 | 0.342 |
| -0.3 | 0.927 | 5.048 | 1.197 | 60.000 | 0.163 | 0.000 | 0.254 |
| -0.4 | 0.360 | 0.602 | 0.217 | 25.500 | 0.107 | 0.000 | 0.638 |
| -0.5 | 0.000 | 0.000 | 0.000 | 0.000 | 0.000 | 0.000 | 0.471 |
| -0.6 | 0.000 | 0.000 | 0.000 | 0.000 | 0.000 | 0.000 | 0.622 |

All data are expressed as median values. Legend: Thr, R threshold; $\lambda$ , $\alpha$ model parameters; $x_{min}$ , lower bound for model distribution; TL, length of the tail; KS, Kolgomorov-Smirnov statistic; $p$ -value, plausibility of the model; TL<sub>r</sub>, proportion of non-zero nodes in the tail.

**Table 28.** Fit results of the power law with exponential cutoff distribution for the 20K resolution dataset.

| Thr | $\alpha$ | $\lambda$ | $x_{min}$ | TL | KS | p-value | TL<sub>r</sub> |
| --- | --- | --- | --- | --- | --- | --- | --- |
| +0.8 | 0.150 | 0.046 | 0.402 | 25.500 | 0.039 | 0.000 | 0.489 |
| +0.7 | 1.767 | 0.058 | 1.481 | 98.500 | 0.128 | 0.130 | 0.443 |
| +0.6 | 0.719 | 0.052 | 2.239 | 181.000 | 0.045 | 0.365 | 0.274 |
| +0.5 | 0.836 | 0.024 | 2.505 | 612.000 | 0.035 | 0.450 | 0.358 |
| +0.4 | 0.887 | 0.009 | 1.848 | 2509.000 | 0.019 | 0.190 | 0.331 |
| +0.3 | 0.807 | 0.005 | 3.518 | 2792.000 | 0.015 | 0.060 | 0.293 |
| +0.2 | 0.673 | 0.003 | 5.534 | 4697.000 | 0.013 | 0.020 | 0.295 |
| +0.1 | 0.105 | 0.003 | 76.096 | 8323.000 | 0.013 | 0.020 | 0.415 |
| +0.0 | 0.001 | 0.004 | 624.430 | 9088.000 | 0.021 | 0.005 | 0.452 |
| -0.0 | 2.487 | 0.002 | 223.250 | 5195.000 | 0.010 | 0.590 | 0.259 |
| -0.1 | 1.275 | 0.013 | 4.567 | 4362.500 | 0.012 | 0.720 | 0.218 |
| -0.2 | 1.780 | 0.008 | 3.296 | 701.000 | 0.032 | 0.475 | 0.236 |
| -0.3 | 2.198 | 0.000 | 1.139 | 164.500 | 0.073 | 0.000 | 0.375 |
| -0.4 | 1.141 | 0.000 | 0.200 | 72.500 | 0.020 | 0.000 | 0.315 |
| -0.5 | 0.000 | 0.000 | 0.000 | 0.000 | 0.000 | 0.000 | 0.482 |
| -0.6 | 0.000 | 0.000 | 0.000 | 0.000 | 0.000 | 0.000 | 0.672 |

All data are expressed as median values. Legend: Thr, R threshold; $b$ , $\alpha$ model parameters; $x_{min}$ , lower bound for model distribution; TL, length of the tail; KS, Kolgomorov-Smirnov statistic; $p$ -value, plausibility of the model; TL<sub>r</sub>, proportion of non-zero nodes in the tail.

**Table 29.**
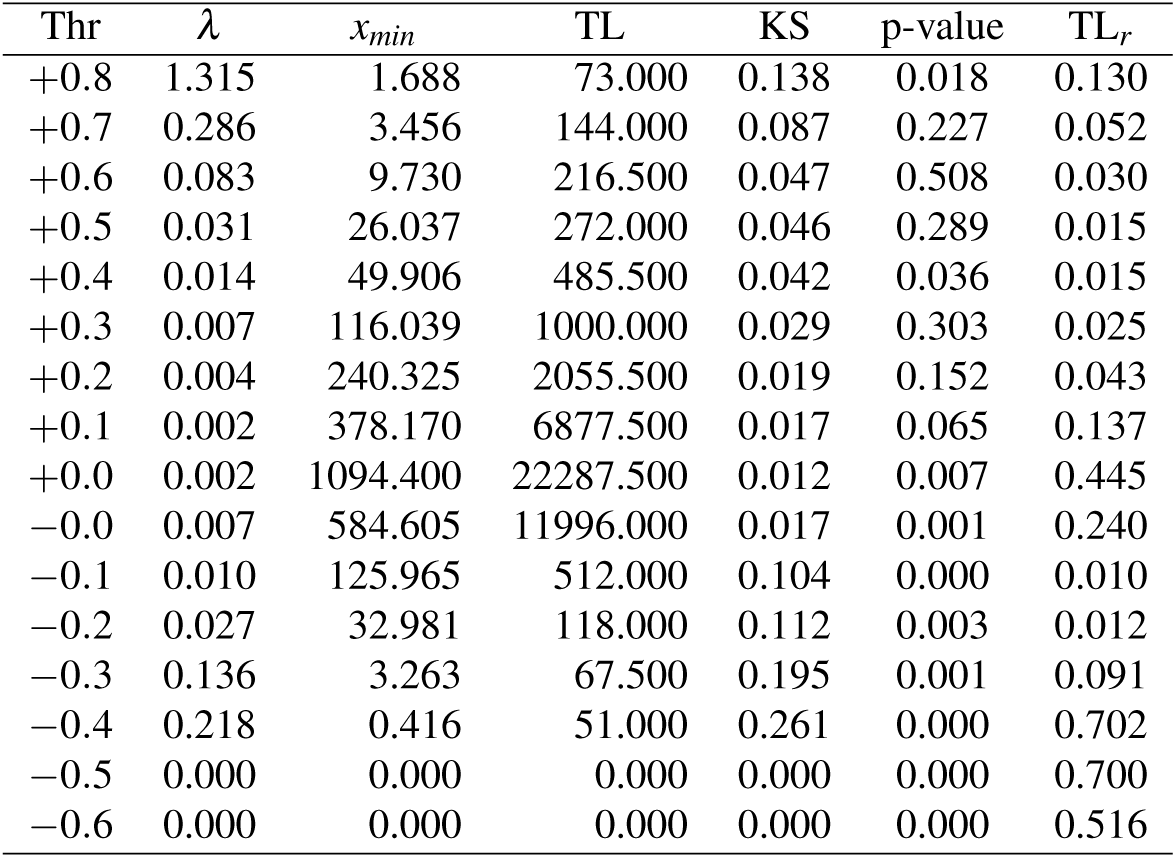
Fit results of the exponential distribution for the 50K resolution dataset.

**Table 30.** Fit results of the power law distribution for the 50K resolution dataset.

| Thr | $\alpha$ | $x_{min}$ | TL | KS | p-value | TL<sub>r</sub> |
| --- | --- | --- | --- | --- | --- | --- |
| +0.8 | 4.878 | 1.662 | 100.000 | 0.143 | 0.002 | 0.141 |
| +0.7 | 3.455 | 1.608 | 265.500 | 0.089 | 0.002 | 0.153 |
| +0.6 | 2.340 | 1.832 | 732.000 | 0.066 | 0.024 | 0.080 |
| +0.5 | 2.888 | 2.623 | 516.500 | 0.066 | 0.001 | 0.038 |
| +0.4 | 2.649 | 2.156 | 4458.500 | 0.068 | 0.000 | 0.140 |
| +0.3 | 2.287 | 0.706 | 24476.500 | 0.063 | 0.000 | 0.594 |
| +0.2 | 1.636 | 1.643 | 23799.000 | 0.063 | 0.000 | 0.496 |
| +0.1 | 5.519 | 1340.500 | 959.000 | 0.069 | 0.000 | 0.019 |
| +0.0 | 5.926 | 1649.000 | 8378.500 | 0.052 | 0.000 | 0.168 |
| -0.0 | 7.738 | 851.895 | 3342.000 | 0.012 | 0.797 | 0.067 |
| -0.1 | 2.382 | 25.005 | 12457.000 | 0.024 | 0.000 | 0.248 |
| -0.2 | 2.259 | 8.079 | 2045.500 | 0.041 | 0.000 | 0.157 |
| -0.3 | 2.306 | 1.322 | 470.000 | 0.057 | 0.018 | 0.218 |
| -0.4 | 2.446 | 0.697 | 59.500 | 0.082 | 0.001 | 0.345 |
| -0.5 | 0.000 | 0.000 | 0.000 | 0.000 | 0.000 | 0.307 |
| -0.6 | 0.000 | 0.000 | 0.0 | 0.000 | 0.0000 | 0.344 |

All data are expressed as median values. Legend: Thr, R threshold; $\lambda$ , $\alpha$ model parameters; $x_{min}$ , lower bound for model distribution; TL, length of the tail; KS, Kolmogorov-Smirnov statistic; p-value, plausibility of the model; TL<sub>r</sub>, proportion of non-zero nodes in the tail.

**Table 31.** Fit results of the generalized Pareto distribution for the 50K resolution dataset.

| Thr | $k$ | $\sigma$ | $x_{min}$ | TL | KS | p-value | $TL_r$ |
| --- | --- | --- | --- | --- | --- | --- | --- |
| +0.8 | 0.472 | 0.084 | 1.691 | 84.000 | 0.089 | 0.066 | 0.131 |
| +0.7 | 0.311 | 2.282 | 3.004 | 279.000 | 0.057 | 0.147 | 0.106 |
| +0.6 | 0.146 | 8.489 | 6.110 | 279.000 | 0.033 | 0.571 | 0.049 |
| +0.5 | 0.270 | 29.789 | 15.580 | 726.000 | 0.026 | 0.614 | 0.039 |
| +0.4 | 0.513 | 35.703 | 23.940 | 2313.500 | 0.020 | 0.191 | 0.074 |
| +0.3 | -0.010 | 128.614 | 117.816 | 1102.500 | 0.022 | 0.347 | 0.028 |
| +0.2 | -0.093 | 308.690 | 336.890 | 1845.500 | 0.019 | 0.347 | 0.039 |
| +0.1 | -0.168 | 536.680 | 884.885 | 2566.500 | 0.014 | 0.512 | 0.051 |
| +0.0 | -0.059 | 515.560 | 845.705 | 26921.000 | 0.008 | 0.001 | 0.537 |
| -0.0 | 0.159 | 133.645 | 684.440 | 6367.000 | 0.007 | 0.378 | 0.127 |
| -0.1 | 0.594 | 26.092 | 12.098 | 19056.000 | 0.012 | 0.000 | 0.380 |
| -0.2 | 0.634 | 4.589 | 1.430 | 2449.000 | 0.019 | 0.224 | 0.295 |
| -0.3 | 0.596 | 2.443 | 0.935 | 434.000 | 0.033 | 0.011 | 0.328 |
| -0.4 | 0.692 | 0.091 | 0.410 | 137.500 | 0.066 | 0.000 | 0.718 |
| -0.5 | 0.000 | 0.000 | 0.000 | 0.0 | 0.000 | 0.0000 | 0.243 |
| -0.6 | 0.000 | 0.000 | 0.000 | 0.0 | 0.000 | 0.0000 | 0.637 |

All data are expressed as median values. Legend: Thr, R threshold; $k$ , $\sigma$ , $\mu$ model parameters; $x_{min}$ , lower bound for model distribution; TL, length of the tail; KS, Kolgomorov-Smirnov statistic; $p$ -value, plausibility of the model; $TL_r$ , proportion of non-zero nodes in the tail.

**Table 32.** Fit results of the log-normal distribution for the 50K resolution dataset.

| Thr | $\mu$ | $\sigma$ | $x_{min}$ | TL | KS | p-value | $TL_r$ |
| --- | --- | --- | --- | --- | --- | --- | --- |
| +0.8 | 0.589 | 0.626 | 1.701 | 75.000 | 0.169 | 0.007 | 0.126 |
| +0.7 | 1.242 | 1.169 | 2.679 | 161.500 | 0.067 | 0.112 | 0.061 |
| +0.6 | 2.418 | 0.926 | 7.344 | 262.000 | 0.038 | 0.400 | 0.035 |
| +0.5 | 2.886 | 0.849 | 6.419 | 329.500 | 0.032 | 0.510 | 0.019 |
| +0.4 | 3.655 | 0.900 | 25.373 | 827.500 | 0.026 | 0.342 | 0.026 |
| +0.3 | 4.561 | 0.843 | 72.916 | 1370.500 | 0.022 | 0.050 | 0.034 |
| +0.2 | 5.570 | 0.586 | 229.450 | 1828.000 | 0.019 | 0.030 | 0.039 |
| +0.1 | 6.854 | 0.405 | 957.705 | 2240.500 | 0.016 | 0.147 | 0.045 |
| +0.0 | 6.768 | 0.493 | 729.410 | 28046.000 | 0.007 | 0.003 | 0.559 |
| -0.0 | 6.027 | 0.418 | 456.595 | 34350.000 | 0.008 | 0.007 | 0.687 |
| -0.1 | 2.334 | 1.345 | 12.872 | 5050.500 | 0.012 | 0.112 | 0.101 |
| -0.2 | 1.050 | 1.538 | 1.103 | 2108.500 | 0.021 | 0.250 | 0.260 |
| -0.3 | -0.458 | 1.478 | 0.741 | 413.500 | 0.042 | 0.012 | 0.416 |
| -0.4 | 0.000 | 0.561 | 0.220 | 96.000 | 0.024 | 0.000 | 0.368 |
| -0.5 | 0.000 | 0.000 | 0.000 | 0.000 | 0.000 | 0.000 | 0.625 |
| -0.6 | 0.000 | 0.000 | 0.000 | 0.000 | 0.000 | 0.000 | 0.559 |

All data are expressed as median values. Legend: Thr, R threshold; $\lambda$ , $\alpha$ model parameters; $x_{min}$ , lower bound for model distribution; TL, length of the tail; KS, Kolgomorov-Smirnov statistic; $p$ -value, plausibility of the model; $TL_r$ , proportion of non-zero nodes in the tail.

**Table 33.** Fit results of the Weibull distribution for the 50K resolution dataset.

| Thr | $b$ | $a$ | $x_{min}$ | TL | KS | p-value | $TL_r$ |
| --- | --- | --- | --- | --- | --- | --- | --- |
| +0.8 | 2.545 | 2.808 | 1.340 | 94.500 | 0.182 | 0.000 | 0.195 |
| +0.7 | 2.102 | 6.799 | 3.095 | 163.000 | 0.118 | 0.000 | 0.086 |
| +0.6 | 1.443 | 20.377 | 5.479 | 754.000 | 0.133 | 0.000 | 0.082 |
| +0.5 | 1.599 | 42.253 | 5.893 | 233.000 | 0.112 | 0.000 | 0.015 |
| +0.4 | 1.066 | 64.165 | 6.527 | 1801.500 | 0.110 | 0.000 | 0.056 |
| +0.3 | 0.869 | 91.490 | 6.296 | 4791.000 | 0.109 | 0.000 | 0.116 |
| +0.2 | 0.780 | 143.668 | 6.806 | 11589.500 | 0.107 | 0.000 | 0.240 |
| +0.1 | 0.780 | 158.921 | 16.408 | 34078.000 | 0.095 | 0.000 | 0.679 |
| +0.0 | 2.362 | 1185.807 | 301.736 | 50112.500 | 0.082 | 0.000 | 1.000 |
| -0.0 | 3.281 | 596.815 | 298.105 | 50052.000 | 0.149 | 0.000 | 1.000 |
| -0.1 | 0.722 | 12.127 | 0.324 | 50100.000 | 0.173 | 1.000 | 1.000 |
| -0.2 | 1.006 | 20.542 | 4.651 | 641.000 | 0.141 | 0.000 | 0.118 |
| -0.3 | 0.973 | 17.591 | 3.652 | 73.500 | 0.158 | 0.000 | 0.058 |
| -0.4 | 0.812 | 1.596 | 0.444 | 51.000 | 0.210 | 0.000 | 0.569 |
| -0.5 | 0.000 | 0.000 | 0.000 | 0.000 | 0.000 | 0.000 | 0.559 |
| -0.6 | 0.000 | 0.000 | 0.000 | 0.000 | 0.000 | 0.000 | 0.518 |

All data are expressed as median values. Legend: Thr, R threshold; $\lambda$ , $\alpha$ model parameters; $x_{min}$ , lower bound for model distribution; TL, length of the tail; KS, Kolgomorov-Smirnov statistic; $p$ -value, plausibility of the model; $TL_r$ , proportion of non-zero nodes in the tail.

**Table 34.** Fit results of the exponential distribution for the 80K resolution dataset.

| Thr | $\lambda$ | $x_{min}$ | TL | KS | p-value | $TL_r$ |
| --- | --- | --- | --- | --- | --- | --- |
| +0.8 | 0.723 | 2.424 | 80.500 | 0.149 | 0.034 | 0.052 |
| +0.7 | 0.225 | 4.907 | 192.500 | 0.065 | 0.269 | 0.038 |
| +0.6 | 0.079 | 10.428 | 436.000 | 0.038 | 0.338 | 0.029 |
| +0.5 | 0.033 | 28.607 | 909.000 | 0.060 | 0.021 | 0.022 |
| +0.4 | 0.013 | 65.146 | 804.500 | 0.047 | 0.007 | 0.016 |
| +0.3 | 0.006 | 134.157 | 1262.500 | 0.031 | 0.034 | 0.018 |
| +0.2 | 0.003 | 345.358 | 2190.500 | 0.019 | 0.150 | 0.029 |
| +0.1 | 0.002 | 513.945 | 7653.000 | 0.017 | 0.017 | 0.092 |
| +0.0 | 0.005 | 933.312 | 19524.500 | 0.015 | 0.000 | 0.256 |
| -0.0 | 0.005 | 933.312 | 19524.500 | 0.015 | 0.000 | 0.256 |
| -0.1 | 0.009 | 139.661 | 902.500 | 0.099 | 0.000 | 0.011 |
| -0.2 | 0.024 | 29.743 | 269.500 | 0.159 | 0.000 | 0.017 |
| -0.3 | 0.071 | 7.720 | 81.500 | 0.153 | 0.009 | 0.051 |
| -0.4 | 0.346 | 0.472 | 55.000 | 0.270 | 0.000 | 0.569 |
| -0.5 | 0.024 | 0.259 | 25.500 | 0.107 | 0.000 | 0.630 |
| -0.6 | 0.000 | 0.000 | 0.000 | 0.000 | 0.000 | 0.505 |

**Table 35.** Fit results of the power law distribution for the 80K resolution dataset.

| Thr | $\alpha$ | $x_{min}$ | TL | KS | p-value | TL<sub>r</sub> |
| --- | --- | --- | --- | --- | --- | --- |
| +0.8 | 3.867 | 2.464 | 72.500 | 0.149 | 0.003 | 0.053 |
| +0.7 | 2.838 | 2.187 | 345.000 | 0.067 | 0.010 | 0.058 |
| +0.6 | 3.056 | 2.143 | 1244.000 | 0.056 | 0.002 | 0.075 |
| +0.5 | 2.413 | 2.441 | 3677.000 | 0.065 | 0.000 | 0.102 |
| +0.4 | 2.741 | 2.134 | 12172.000 | 0.056 | 0.000 | 0.190 |
| +0.3 | 2.525 | 2.746 | 21472.000 | 0.063 | 0.000 | 0.292 |
| +0.2 | 1.717 | 4.314 | 11952.500 | 0.065 | 0.000 | 0.151 |
| +0.1 | 2.707 | 694.927 | 37578.500 | 0.059 | 0.000 | 0.468 |
| +0.0 | 8.273 | 1258.367 | 5745.500 | 0.011 | 0.362 | 0.075 |
| -0.0 | 8.273 | 1258.367 | 5745.500 | 0.011 | 0.354 | 0.075 |
| -0.1 | 2.312 | 19.290 | 13168.500 | 0.023 | 0.000 | 0.170 |
| -0.2 | 2.237 | 11.680 | 2844.000 | 0.035 | 0.000 | 0.132 |
| -0.3 | 2.240 | 2.666 | 681.000 | 0.054 | 0.042 | 0.223 |
| -0.4 | 2.375 | 1.107 | 111.000 | 0.107 | 0.000 | 0.315 |
| -0.5 | 1.068 | 0.266 | 25.500 | 0.019 | 0.000 | 0.293 |
| -0.6 | 0.000 | 0.000 | 0.000 | 0.000 | 0.000 | 0.410 |

All data are expressed as median values. Legend: Thr, R threshold; $\lambda$ , $\alpha$ model parameters; $x_{min}$ , lower bound for model distribution; TL, length of the tail; KS, Kolgomorov-Smirnov statistic; $p$ -value, plausibility of the model; TL<sub>r</sub>, proportion of non-zero nodes in the tail.

**Table 36.** Fit results of the generalized Pareto distribution for the 80K resolution dataset.

| Thr | $k$ | $\sigma$ | $x_{min}$ | TL | KS | p-value | TL<sub>r</sub> |
| --- | --- | --- | --- | --- | --- | --- | --- |
| +0.8 | 0.698 | 0.163 | 2.107 | 130.000 | 0.096 | 0.004 | 0.118 |
| +0.7 | 0.297 | 2.578 | 3.675 | 190.500 | 0.044 | 0.185 | 0.042 |
| +0.6 | 0.119 | 7.267 | 6.109 | 570.000 | 0.025 | 0.532 | 0.031 |
| +0.5 | 0.306 | 18.800 | 14.445 | 961.000 | 0.019 | 0.335 | 0.030 |
| +0.4 | 0.303 | 48.210 | 33.473 | 1800.000 | 0.021 | 0.120 | 0.034 |
| +0.3 | 0.154 | 106.347 | 60.200 | 2383.500 | 0.022 | 0.195 | 0.035 |
| +0.2 | -0.075 | 363.671 | 468.904 | 1734.500 | 0.018 | 0.400 | 0.022 |
| +0.1 | -0.144 | 655.662 | 1432.557 | 2416.500 | 0.010 | 0.767 | 0.031 |
| +0.0 | 0.187 | 179.022 | 1139.975 | 5933.000 | 0.009 | 0.147 | 0.080 |
| -0.0 | 0.187 | 179.022 | 1139.975 | 5933.000 | 0.009 | 0.140 | 0.080 |
| -0.1 | 0.590 | 39.240 | 23.752 | 23584.000 | 0.011 | 0.000 | 0.289 |
| -0.2 | 0.664 | 6.544 | 3.499 | 2992.500 | 0.016 | 0.253 | 0.211 |
| -0.3 | 0.663 | 2.680 | 1.164 | 389.000 | 0.030 | 0.292 | 0.312 |
| -0.4 | 0.707 | 0.747 | 0.858 | 86.500 | 0.059 | 0.003 | 0.464 |
| -0.5 | 0.232 | 0.025 | 0.250 | 33.000 | 0.018 | 0.000 | 0.257 |
| -0.6 | 0.000 | 0.000 | 0.000 | 0.000 | 0.000 | 0.000 | 0.564 |
| -0.7 | 0.000 | 0.000 | 0.000 | 0.000 | 0.000 | 0.000 | 0.773 |

All data are expressed as median values. Legend: Thr, R threshold; $k$ , $\sigma$ , $\mu$ model parameters; $x_{min}$ , lower bound for model distribution; TL, length of the tail; KS, Kolgomorov-Smirnov statistic; $p$ -value, plausibility of the model; TL<sub>r</sub>, proportion of non-zero nodes in the tail.

**Table 37.** Fit results of the Weibull distribution for the 80K resolution dataset.

| Thr | $b$ | $a$ | $x_{min}$ | TL | KS | p-value | TL<sub>r</sub> |
| --- | --- | --- | --- | --- | --- | --- | --- |
| +0.8 | 2.354 | 3.535 | 1.841 | 100.000 | 0.168 | 0.000 | 0.087 |
| +0.7 | 1.950 | 9.263 | 3.790 | 204.000 | 0.142 | 0.000 | 0.047 |
| +0.6 | 1.396 | 20.110 | 5.254 | 653.500 | 0.139 | 0.000 | 0.044 |
| +0.5 | 1.125 | 36.690 | 6.423 | 2215.500 | 0.140 | 0.000 | 0.056 |
| +0.4 | 1.050 | 86.351 | 7.985 | 2232.000 | 0.122 | 0.000 | 0.042 |
| +0.3 | 0.930 | 137.916 | 10.353 | 5415.500 | 0.116 | 0.000 | 0.077 |
| +0.2 | 0.808 | 191.619 | 9.965 | 14371.500 | 0.117 | 0.000 | 0.186 |
| +0.1 | 0.781 | 322.665 | 27.748 | 46635.000 | 0.118 | 0.000 | 0.645 |
| +0.0 | 3.715 | 960.365 | 529.864 | 76752.500 | 0.153 | 0.000 | 1.000 |
| -0.0 | 3.715 | 960.365 | 529.864 | 76752.500 | 0.153 | 0.000 | 1.000 |
| -0.1 | 0.885 | 35.413 | 1.006 | 70190.000 | 0.187 | 0.396 | 1.000 |
| -0.2 | 0.704 | 9.017 | 0.579 | 6616.500 | 0.150 | 0.801 | 0.447 |
| -0.3 | 0.982 | 17.408 | 5.343 | 98.000 | 0.151 | 0.000 | 0.051 |
| -0.4 | 0.909 | 2.253 | 0.455 | 51.500 | 0.205 | 0.000 | 0.479 |
| -0.5 | 0.357 | 0.797 | 0.258 | 25.500 | 0.087 | 0.001 | 0.562 |
| -0.6 | 0.000 | 0.000 | 0.000 | 0.000 | 0.000 | 0.000 | 0.550 |

All data are expressed as median values. Legend: Thr, R threshold; $\lambda$ , $\alpha$ model parameters; $x_{min}$ , lower bound for model distribution; TL, length of the tail; KS, Kolmogorov-Smirnov statistic; p-value, plausibility of the model; TL<sub>r</sub>, proportion of non-zero nodes in the tail.

**Table 38.** Likelihood ratio test results from comparing the best fit for alternative distributions with the best fit power law distribution for the 1K dataset. We show the percentage of times a power law model (M*_PL_*), the alternative model (M*_Alt_*) or neither was favored.

| Alternative | $M_{PL}$ | $M_{Alt}$ | Inconclusive |
| --- | --- | --- | --- |
| Exponential | 22.078 (17/77) | 40.260 (31/77) | 37.662 (29/77) |
| Truncated PL | 83.333 (55/66) | 6.061 (4/66) | 10.606 (7/66) |
| Log normal | 3.261 (3/92) | 41.304 (38/92) | 55.435 (51/92) |
| Weibull | 93.939 (62/66) | 6.061 (4/66) | 0.000 (0/66) |
| Generalized Pareto | 4.040 (4/99) | 45.455 (45/99) | 50.505 (50/99) |

**Table 39.** Likelihood ratio test results from comparing the best fit for alternative distributions with the best fit power law distribution for the 5K dataset. We show the percentage of times a power law model (M*_PL_*), the alternative model (M*_Alt_*) or neither was favored.

| Alternative | $M_{PL}$ | $M_{Alt}$ | Inconclusive |
| --- | --- | --- | --- |
| Exponential | 25.806 (24/93) | 51.613 (48/93) | 22.581 (21/93) |
| Truncated PL | 69.014 (49/71) | 19.718 (14/71) | 11.268 (8/71) |
| Log normal | 9.091 (9/99) | 50.505 (50/99) | 40.404 (40/99) |
| Weibull | 81.690 (58/71) | 18.310 (13/71) | 0.000 (0/71) |
| Generalized Pareto | 7.767 (8/103) | 51.456 (53/103) | 40.777 (42/103) |

**Table 40.** Likelihood ratio test results from comparing the best fit for alternative distributions with the best fit power law distribution for the 10K dataset. We show the percentage of times a power law model (M*_PL_*), the alternative model (M*_Alt_*) or neither was favored.

| Alternative | $M_{PL}$ | $M_{Alt}$ | Inconclusive |
| --- | --- | --- | --- |
| Exponential | 29.070 (25/86) | 58.140 (50/86) | 12.791 (11/86) |
| Truncated PL | 49.254 (33/67) | 34.328 (23/67) | 16.418 (11/67) |
| Log normal | 9.574 (9/94) | 61.702 (58/94) | 28.723 (27/94) |
| Weibull | 52.239 (35/67) | 34.328 (23/67) | 13.433 (9/67) |
| Generalized Pareto | 2.970 (3/101) | 64.356 (65/101) | 32.673 (33/101) |

**Table 41.** Likelihood ratio test results from comparing the best fit for alternative distributions with the best fit power law distribution for the 20K dataset. We show the percentage of times a power law model (M*_PL_*), the alternative model (M*_Alt_*) or neither was favored.

| Alternative | $M_{PL}$ | $M_{Alt}$ | Inconclusive |
| --- | --- | --- | --- |
| Exponential | 30.000 (24/80) | 61.250 (49/80) | 8.750 (7/80) |
| Truncated PL | 50.000 (27/54) | 35.185 (19/54) | 14.815 (8/54) |
| Log normal | 9.639 (8/83) | 62.651 (52/83) | 27.711 (23/83) |
| Weibull | 57.407 (31/54) | 35.185 (19/54) | 7.407 (4/54) |
| Generalized Pareto | 6.0 (6/100) | 72.0 (72/100) | 22.0 (22/100) |

**Table 42.**
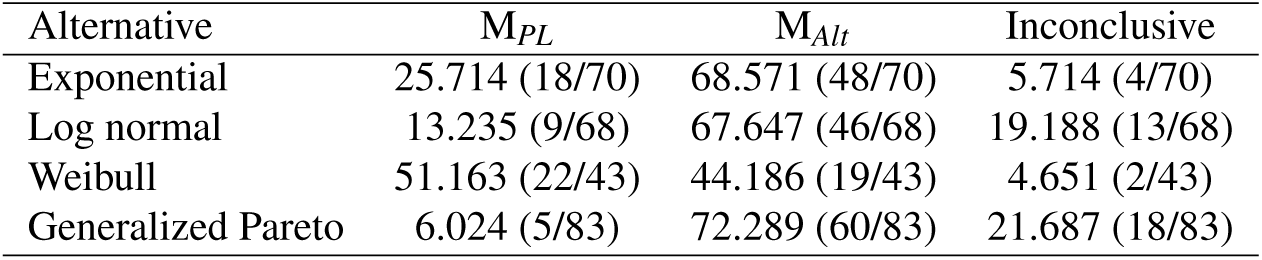
Likelihood ratio test results from comparing the best fit for alternative distributions with the best fit power law distribution for the 50K dataset. We show the percentage of times a power law model (M*_PL_*), the alternative model (M*_Alt_*) or neither was favored.

| Alternative | $M_{PL}$ | $M_{Alt}$ | Inconclusive |
| --- | --- | --- | --- |
| Exponential | 25.714 (18/70) | 68.571 (48/70) | 5.714 (4/70) |
| Log normal | 13.235 (9/68) | 67.647 (46/68) | 19.188 (13/68) |
| Weibull | 51.163 (22/43) | 44.186 (19/43) | 4.651 (2/43) |
| Generalized Pareto | 6.024 (5/83) | 72.289 (60/83) | 21.687 (18/83) |

**Table 43.** Likelihood ratio test results from comparing the best fit for alternative distributions with the best fit power law distribution for the 80K dataset. We show the percentage of times a power law model (M*_PL_*), the alternative model (M*_Alt_*) or neither was favored.

| Alternative | $M_{PL}$ | $M_{Alt}$ | Inconclusive |
| --- | --- | --- | --- |
| Exponential | 39.062 (25/64) | 54.688 (35/64) | 6.250 (4/64) |
| Weibull | 62.500 (30/48) | 37.500 (18/48) | 0.000 (0/48) |
| Generalized Pareto | 6.977 (6/86) | 68.605 (59/86) | 24.419 (21/86) |

